# The non-steroidal MR antagonist Finerenone reverses Western diet-induced kidney disease by regulating mitochondrial and lipid metabolism and inflammation

**DOI:** 10.1101/2025.04.02.646892

**Authors:** Komuraiah Myakala, Xiaoxin X. Wang, Nataliia Shults, Eleni P. Hughes, Patricia de Carvalho Ribeiro, Rozhin Penjweini, Katie Link, Keely Barton, Ewa Krawczyk, Cheryl Clarkson Paredes, Anastas Popratiloff, Jay R. Knutson, Ashley L Cowart, Moshe Levi

## Abstract

Mineralocorticoid receptor (MR) overactivation plays a crucial role in the pathogenesis of chronic kidney disease, as well as several cardiovascular and arterial diseases. Current studies determined the mechanisms of the beneficial kidney effects of the non-steroidal MR antagonist Finerenone (FN) in a mouse model of western diet-induced obesity and insulin resistance. 10-week-old male C57BL/6J mice were fed a low fat (LF) or a western diet (WD) for 12 weeks followed by treatment with either vehicle or finerenone (FN) for another 14 weeks (intervention studies) until they were 36 weeks old. Finerenone treatment prevented a) the increased albuminuria and kidney injury molecule 1 (KIM1), b) the expanded extracellular mesangial matrix, and podocyte injury, c) fibronectin, collagen IV, CD45 and CD68 immunostaining, d) glomerular basement membrane disruption, podocyte foot process loss, and mitochondrial structural abnormalities, e) the pro-inflammatory cytokines (MCP1), innate immunity pathways (TLR2, STING, STAT3), and fibrosis markers fibronectin, TGFβ and Pai1, and f) the increased kidney cholesterol levels. There was also reduced expression of nuclear receptor ERRγ without changes in ERRα in WD-fed mice whereas both ERRα and ERRγ expression levels increased after Finerenone treatment. NADH lifetime analysis showed decreased bound NADH, compatible with decreased mitochondrial OXPHOS in the kidneys of WD-fed mice compared to controls, which was prevented by finerenone treatment. In conclusion, Finerenone treatment exhibits a renal protective role and prevents the progression of kidney disease by regulating mitochondrial function, most likely via ERRγ, and reducing lipid accumulation and inflammation.

## INTRODUCTION

Chronic kidney disease (CKD) affects approximately 850 million people worldwide, with 3.9 million requiring kidney replacement (1). Up to 40% of people with diabetes develop CKD, making diabetic kidney disease (DKD) the leading cause of kidney failure globally (2). The increasing prevalence of type 2 diabetes (T2D) has significantly raised the rate of CKD diagnoses. Individuals diagnosed with both conditions face a substantially higher risk of mortality, often experiencing a three-fold increase compared to those with only T2D (3, 4). Additionally, cardiovascular mortality in CKD stage 3 exceeds the risk of kidney failure by tenfold, positioning CKD as the fifth leading cause of death globally by 2040, primarily due to its association with T2D (5–7).

Several factors, such as obesity, hypertension, and insulin resistance, are recognized as critical risk determinants for chronic kidney disease (CKD) and cardiovascular complications, even without diabetes or hyperglycemia (8–11). These conditions promote systemic inflammation, oxidative stress, and endothelial dysfunction, driving renal and cardiovascular damage (9, 11–13). Angiotensin-converting enzyme inhibitors (ACE inhibitors) and angiotensin receptor blockers (ARBs) have been important in managing albuminuric CKD for over two decades (14, 15). These agents mitigate the progression of CKD by inhibiting the renin-angiotensin system, thereby reducing intraglomerular pressure and proteinuria. Their efficacy in slowing CKD progression and reducing cardiovascular events has been documented in landmark trials (14–16).

While renin-angiotensin system (RAS) blockers such as angiotensin-converting enzyme inhibitors (ACEis) and angiotensin receptor blockers (ARBs) offer significant kidney protection, a substantial risk of chronic kidney disease (CKD) and cardiovascular disease progression persists. Attempts to enhance therapeutic outcomes through dual RAS blockade combining ACEis, ARBs, or direct renin inhibitors have been largely unsuccessful due to increased risks of adverse events (17–20). Like ACEis and ARBs, SGLT2 inhibitors are another class of medications that have demonstrated significant benefits in managing CKD, both in patients with and without diabetes. Clinical trials, such as EMPA-KIDNEY, and seminal randomized controlled trials like CREDENCE and DAPA-CKD, have highlighted their efficacy in slowing eGFR decline, reducing albuminuria, and improving cardiovascular and renal outcomes (21). GLP-1 receptor agonists have also been associated with lower all-cause mortality and lower sepsis- and infection-related mortality in patients with diabetes and advanced CKD or ESKD (22).

Interestingly, the mineralocorticoid receptor (MR) plays a key role in the aldosterone-sensitive distal nephron, regulating sodium (Na⁺) and potassium (K⁺) homeostasis, extracellular fluid volume, and blood pressure (23–27). Aldosterone secretion is primarily stimulated by elevated blood K⁺ levels or activation of the renin-angiotensin-aldosterone system (RAAS) due to reduced blood volume or sodium intake. By forming a complex with MR in principal distal nephron cells, aldosterone enhances Na⁺ reabsorption and K⁺ excretion through the activation of epithelial sodium channels (ENaC) and Na⁺–K⁺ ATPase (28, 29). In intercalated cells, the aldosterone-MR complex upregulates pendrin to increase chloride (Cl⁻) reabsorption and decrease K⁺ secretion, while in distal convoluted tubule cells, it boosts Na⁺ and Cl⁻ reabsorption by enhancing sodium–chloride cotransporter (NCC) activity. Overactivation of the MR in animal models of chronic kidney disease (CKD) has been shown to exacerbate sodium retention and hypertension, while simultaneously driving pathological processes such as inflammation and fibrosis within the kidneys, vasculature, and heart (24, 30–32). These maladaptive responses contribute significantly to the progression of cardiorenal disease, a condition characterized by the interplay of kidney and cardiovascular dysfunction. The resulting tissue damage and systemic effects underscore the pivotal role of MR signaling in disease pathogenesis. Therapeutically, targeting MR overactivation presents a compelling strategy to mitigate CKD progression and its associated cardiovascular complications (26, 27, 33, 34). MR antagonism not only reduces sodium retention and blood pressure but also attenuates inflammatory and fibrotic pathways, offering dual benefits in preserving renal function and improving cardiovascular outcomes. By addressing both renal and cardiac damage, MR blockade holds the potential to significantly lower morbidity and mortality in patients with CKD, making it a critical focus in the development of novel cardiorenal therapies (26, 27, 33, 34). The pressing need for therapies to effectively slow the progression of cardiorenal disease with improved safety profiles has driven substantial efforts toward the development of nonsteroidal mineralocorticoid receptor antagonists (MRAs) (25–27, 35–40). Recent development and meta-analysis studies demonstrated that the combination of MRAs and SGLT2 inhibitors resulted in greater reductions in albuminuria and blood pressure compared to SGLT2 inhibitors alone, clearly demonstrating the promising role of MRAs in providing renal benefits (41). Finerenone, a novel nonsteroidal MRA, stands out due to its unique binding characteristics, which confer high potency and selectivity for the MR. Preclinical studies have demonstrated that finerenone exerts anti-inflammatory, antifibrotic, and antiproliferative effects, contributing to the modulation of pathological tissue remodeling in both the heart and kidneys (29, 31, 38, 39, 42–46). Clinical evidence supports finerenone’s efficacy and safety. Phase II trials reported dose-dependent reductions in albuminuria with a safety profile comparable to placebo, while also showing smaller increases in serum potassium levels relative to spironolactone (29). Furthermore, in a pivotal Phase III trial involving patients with CKD and type 2 diabetes mellitus (T2DM), finerenone significantly improved long-term kidney and cardiovascular outcomes compared to placebo (44). These findings highlight finerenone’s potential as a transformative therapy for managing CKD and associated cardiovascular risks, addressing a critical unmet need in this patient population.

Obesity is a well-established and potent risk factor for the development and progression of kidney disease. It significantly elevates the risk of major contributing conditions to chronic kidney disease (CKD), including diabetes mellitus and hypertension, both of which are leading causes of CKD worldwide. Additionally, obesity exerts direct deleterious effects on renal structure and function, independent of these comorbidities. Western diets containing high amounts of saturated fat are major contributors to obesity, acting as metabolic stressors that further aggravate CKD. In the context of diabetic kidney disease (DKD), the Western-type diet worsens metabolic stress by promoting insulin resistance, hyperglycemia, and lipid accumulation, further aggravating DKD progression (47, 48). These factors disrupt the glomerular filtration barrier, leading to proteinuria, a key marker and driver of DKD. Together, these mechanisms accelerate DKD progression and heighten cardiovascular and renal risks.

The renal estrogen-related receptor gamma (ERRγ) pathway is a critical regulator of kidney function, particularly in the context of energy metabolism, oxidative stress, and fibrotic responses. ERRγ plays key roles in the transcriptional regulation of genes involved in mitochondrial function, lipid metabolism, and cellular homeostasis.

While mineralocorticoid receptor (MR) antagonists have shown promise in managing cardiorenal diseases, the specific efficacy and mechanisms of finerenone under conditions of a western diet-induced obesity and insulin resistance remain insufficiently understood.

## MATERIALS AND METHODS

### Animal study

Ten-week-old male C57BL/6J mice were procured from the Jackson Laboratory and housed under standard conditions with a 12-hour light/12-hour dark cycle. Mice were fed either a western diet (WD: 42% milkfat, 34% sucrose, and 0.2% cholesterol; Research Diet, D12079B) or a matched low-fat control diet (LFD: 10 kcal % fat; Research Diet, D14042701) for a duration of 12 weeks. Following this period, the low-fat and western diet-fed mice were randomly assigned to one of four groups: (1) low-fat diet (LF), (2) low-fat diet supplemented with 10 mg/kg of finerenone (LF + FN), (3) Western diet (WD), and (4) western diet supplemented with 10 mg/kg of finerenone (WD + FN). These dietary regimens were maintained for an additional 14 weeks. Food intake and body weight were monitored biweekly throughout the study.

Before the termination of the study, mice were placed in metabolic cages for 24-hour urine collection. Urine samples were analyzed for volume, creatinine, albumin, kidney injury molecule-1 (KIM-1), neutrophil gelatinase-associated lipocalin (NGAL), and thiobarbituric acid reactive substances (TBARS) as previously described (49, 50).

At the end of the study, mice were euthanized via CO₂ inhalation, and blood samples were collected by cardiac puncture for the measurement of glucose, triglycerides, and cholesterol levels. Kidneys were harvested immediately and processed as follows: (a) snap-frozen for RNA extraction, western blotting, and lipid analysis; (b) fixed in 10% formalin for histological evaluation; and (c) embedded in OCT for immunofluorescence microscopy as per standard lab protocols (51, 52). Animal experiments were approved by the Institutional Animal Care and Use Committees at the Department of Comparative Medicine and the Institutional Animal Care and Use Committee (IACUC) at Georgetown University and followed the standards set by National Institute of Health (NIH) guidelines.

### Biochemical Analysis

Fasting blood glucose levels were measured using a glucometer (Elite XL; Bayer, Tarrytown, NY, USA). Plasma potassium (K^+^), triglycerides and cholesterol levels were quantified with commercially available kits (Catalog #NBP3-25780, Catalog #23666410 and #23666202, Novus Biologicals and Pointe Scientific, USA). Urine albumin and creatinine concentrations were assessed using kits from Exocell (Philadelphia, PA, USA) and BioAssay Systems (Hayward, CA, USA), respectively. Urinary thiobarbituric acid reactive substances (TBARS) were measured using the Quantichrom TBARS Assay Kit (BioAssay Systems, Hayward, CA, USA). Urinary kidney injury molecule-1 (KIM-1; Catalog #ab213477) and neutrophil gelatinase-associated lipocalin (NGAL; Catalog #ab199083) concentrations were determined using mouse enzyme-linked immunosorbent assay (ELISA) kits (Abcam, Cambridge, MA, USA). All the assays were done according to the manufacturer’s instructions (51, 52).

### Blood Pressure Measurement

Systolic blood pressure (SBP) was measured using the tail-cuff method (Model BP-2000-M-6, Visitech Systems, Apex, NC, USA) as previously described (53). Briefly, mice were acclimated to the procedure over two days before data collection. Measurements were recorded over several cycles, and mean SBP values were calculated from at least 25–30 measurements per animal at each time point.

### Real-Time Quantitative PCR (qPCR)

Total RNA was isolated from snap-frozen kidney tissue using the NucleoSpin RNA kit (Takara Bio, Germany). Complementary DNA (cDNA) was synthesized using a high-capacity cDNA reverse transcription kit (Applied Biosystems, USA) following the manufacturer’s instructions. Quantitative real-time PCR (qPCR) was performed as previously described, with target gene expression normalized to 18S rRNA levels (51, 52). Primer sequences are provided in Supplemental Table 1.

### Immunoblotting

Kidney tissues (25 mg) were homogenized in T-PER buffer (Thermo Scientific, Catalog #78510) supplemented with protease and phosphatase inhibitors. Total protein concentration was determined using the BCA assay (Thermo Fisher Scientific). Equal amounts of protein were separated on SDS-PAGE gels, transferred onto PVDF membranes, and probed with primary antibodies including fibronectin (Catalog #SAB5700724, Sigma; 1:1000 dilution) and β-actin (Catalog #A5316, Sigma-Aldrich; 1:5000 dilution). Immune complexes were detected with HRP-conjugated secondary antibodies and visualized using the Azure C300 digital imager (Azure Biosystems, Dublin, CA, USA). We randomly pooled every two samples together with in each group, so that we had 4 samples for each group to run the western blot. Densitometry was performed using Fiji ImageJ software (51, 52).

### Histology Staining

Paraffin-embedded kidney tissue sections (5 µm thick) were stained with Periodic Acid-Schiff (PAS) and Picrosirius Red (PSR) following standard protocols performed by the shared resources core facility at Georgetown University (51, 52). Images were captured using a high-capacity digital slide scanner (Aperio ScanScope GT450, Leica Microsystems, Germany). PSR-stained slides were visualized under polarized light and imaged using an IX83 Inverted Microscope (Olympus, Tokyo, Japan). The glomerular mesangial expansion was quantified as the PAS-positive, nucleus-free mesangial area using the ScanScope image analyzer. PSR images were analyzed in Fiji ImageJ, and data were presented as the percentage of positive-stained pixels.

### Immunofluorescence Microscopy

Frozen kidney sections (5 µm thick) were used for immunostaining. Sections were stained with antibodies against fibronectin (SAB5700724, Sigma), type IV collagen (SAB4500375, Sigma), synaptopodin (SAB3500586, Sigma), CD45 (65087-1-Ig, Proteintech) and CD68 (MCA1957, BioRad). Imaging was performed using a confocal microscope (Leica SP8, Germany). Quantification of the positive staining area in the glomerular region was conducted in Fiji ImageJ, as described previously (51, 52).

### Liquid Chromatography–Mass Spectrometry (LC-MS)

Approximately 20–25 mg of frozen kidney tissue was homogenized in methanol and extracted using methanol: chloroform mixture (2:1 ratio) with 10 µL of an internal sphingolipid standard. Samples were incubated overnight and centrifuged. The supernatant evaporated, and the lipid residue was resuspended in methanol: water (1:1 ratio) before injection. Samples were analyzed using the AB Sciex 5500 QTrap system (GenTech Scientific). Relative ceramide levels were quantified using high-performance liquid chromatography-tandem mass spectrometry (HPLC-MS/MS) and normalized to tissue weight.

### Electron Microscopy

Renal cortex tissues were fixed in the 2.5% glutaraldehyde/2%paraformaldehyde/0.05 mol/L cacodylate solution, post-fixed with 1% osmium tetroxide, and embedded in EmBed812. For *transmission electron microscopy* (TEM) imaging acquisition, ultrathin sections (70 nm) were poststained with uranyl acetate and lead citrate and examined in the Talos F200X FEG TEM (FEI, Hillsboro, OR) at 80 kV. Digital electron micrographs were recorded with the TIA software (FEI). For *scanning electron microscopy (SEM)* evaluation, ultrathin sections (120 nm) were mounted in silicon wafers and observed with a Teneo LV FEG SEM (FEI, ThermoFischer Scientific). For optimal results, we used the optiplan mode (high-resolution) equipped with an in-lens T1 detector (Segmented A+B, working distance of 8 mm). Low-magnification images (600×) were first taken for the observation then we performed high magnification tile images of our regions of interest (35,000) using 2 kV and 0.4 current landing voltage.

Glomerular basement membrane (GBM) thickness was measured using calibrated image analysis software (ImageJ). Scanning electron microscopy (SEM) images were acquired at ×35,000 magnification. The pixel-to-nanometer ratio was calibrated using the scale bar present on each image. GBM thickness was measured orthogonally to the capillary loop at multiple points per glomerulus, carefully avoiding oblique sections and regions with artifacts. For each capillary loop, 20 measurements were taken at regular intervals along the capillary wall. Measurements were collected from five glomerular capillary loops per glomerulus, resulting in 50 measurements per glomerulus (54). This sampling strategy provided a robust dataset for statistical analysis. To minimize the influence of occasional overestimations caused by oblique sections, the harmonic mean of all measured values was calculated for each glomerulus. This approach more accurately reflects the true membrane thickness across the capillary circumference, in accordance with stereological principles proposed by Weibel and Gomez (54). A total of ten glomeruli per animal and three animals per group were analyzed. All image analysis was performed under blinded conditions. Foot process width (nm) calculated as (π/4) × (GBM length / number of PFPs) and assessed effacement of podocyte foot process (PFPs) by identifying broadened, flattened, and fused processes with loss of slit diaphragms (55).

The thickness of the glycocalyx was defined as the perpendicular distance from the outer leaflet of the endothelial plasma membrane to the outer margin of the electron-dense glycocalyx layer projecting into the capillary lumen.

In SEM images, the volume fraction of mitochondria was determined using the morphometric technique with a dot grid. Severity of mitochondrial damage was quantified by assessing abnormal mitochondria with irregular shape and size, signs of hypoplasia, inner and outer membrane disruption. Also, swollen cristae electron-lucent matrix, uneven thickening, homogenization, fragmentation together with cristae lose longitudinal orientation, tightness, and regular spacing. The size of each mitochondrion was calculated by using the FIJI ImageJ software (56). Morphometric analysis was performed under blinded conditions by systematic uniform random sampling with the Fiji Software using 20 randomly selected images.

### Fluorescence Lifetime Imaging Microscopy (FLIM)

Two-photon FLIM was performed using an Olympus IX81/FV1000 confocal laser scanning microscope equipped with a Mai Tai BB DeepSee femtosecond laser (Spectra-Physics, Santa Clara, CA, USA). NADH in kidney sections were excited with a wavelength of 750 nm, and emission was detected with a HPM-100-06 Hybrid detector (Becker & Hickl GmbH, Berlin, Germany) with band pass filter 460/60 nm. Time-correlated single-photon counting (TCSPC) histograms were constructed, and FLIM images were analyzed using SPC Image 8.8 software (Becker & Hickl GmbH) (57). Statistical significance was defined as p < 0.05.

### Mitochondrial Complex Activity Assay

Mitochondrial fractions were isolated from frozen kidney tissue as described previously (58). Complex I (NADH dehydrogenase; Catalog # ab109729) and complex IV (cytochrome c oxidase; Catalog # ab109911) activities were measured using assay kits from Abcam (USA) according to the manufacturer’s protocols.

### Statistical Analysis

Data are presented as mean ± standard error. Comparisons between groups were analyzed using a one-way analysis of variance (ANOVA) followed by a Student-Newman-Keuls post hoc test (GraphPad Prism 8, GraphPad Software, Inc., La Jolla, CA, USA). A p-value of <0.05 was considered statistically significant.

## RESULTS

### Effects of finerenone treatment on metabolic and biochemical characteristics in western diet fed mice

Ten-week-old male C57BL/6J mice were fed either a low-fat (LF) or a Western diet (WD) for 12 weeks. Following this period, the mice were maintained on their respective diets, with or without finerenone supplementation, for an additional 14 weeks. Food intake and body weight were recorded every two weeks throughout the study (Supplementary Fig. 1). WD-fed mice consumed more food than LF-fed mice, while finerenone supplementation did not affect food intake compared to diet-matched controls (Supplementary Fig. 1). Consistent with the increased caloric intake, WD-fed mice showed significant weight gain compared to LF-fed controls. However, body weight in either dietary group was unaffected by finerenone treatment. Fasting blood glucose and plasma K^+^ levels were similar among the four groups. Plasma triglyceride and cholesterol levels, which were markedly elevated in WD-fed mice at 36 weeks of age, were significantly reduced with finerenone treatment (Table 1). Systolic blood pressure did not differ significantly across any of the groups.

### Finerenone treatment protects the renal injury in a western diet-induced obesity mouse model

At the end of the finerenone treatment, 24-hour urine samples were collected to assess glomerular filtration barrier function and tubular injury. Urinary albumin levels served as an indicator of glomerular injury, while kidney injury molecule-1 (KIM1) and lipocalin-2 (NGAL) were measured to evaluate tubular damage. Western diet (WD)-fed mice showed a significant increase in urinary albumin, KIM1/creatinine ratio, and NGAL compared to low-fat (LF) fed controls. However, finerenone supplementation significantly reduced these levels in WD-fed mice (Fig. 1A-B). Additionally, WD-fed mice demonstrated increased thiobarbituric acid reactive substances (TBARS), a marker of lipid peroxidation, which was significantly lowered following finerenone treatment (Fig. 1C).

**Figure 1:**
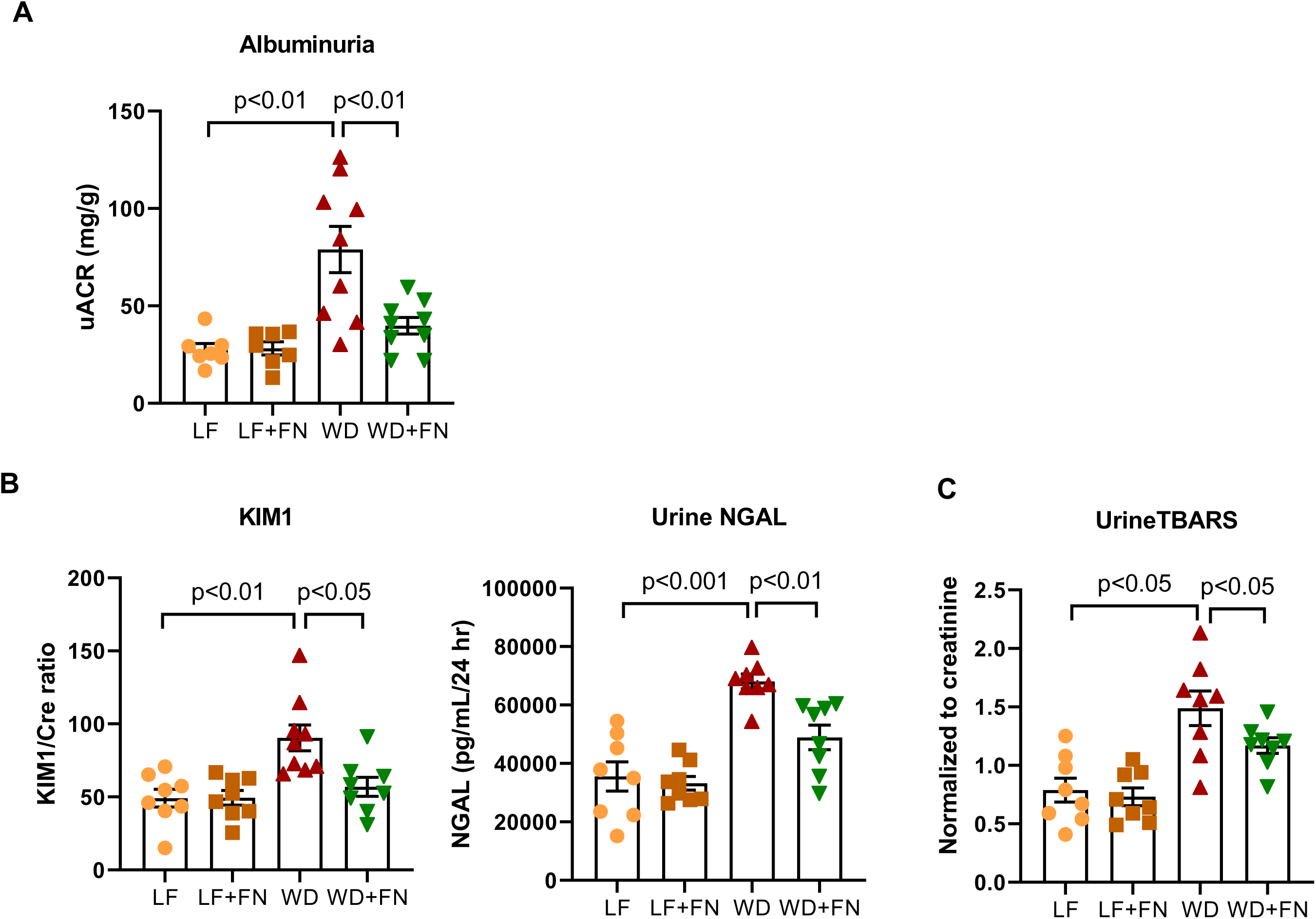
Finerenone treatment improved urine parameters in diet-induced obesity. 24-hour urine measured for albumin, kidney injury molecule 1 (KIM1) normalized to creatinine, neutrophil gelatinase-associated lipocalin (NGAL), and thiobarbituric acid reactive substances (TBARS). n=7-9 mice per group, p-value indicated as p<0.01. uACR, albumin:creatinine ratio.

Diet-induced obesity commonly leads to kidney pathology characterized by increased mesangial matrix accumulation, foot process effacement, and thickening of the glomerular basement membrane (GBM), culminating in renal fibrosis (59). To evaluate how WD-induced obesity and insulin resistance affect renal histopathology, periodic acid Schiff (PAS) and picrosirius red (PSR) staining were performed on kidney sections. In PAS-stained tissues, WD-fed mice exhibited pronounced mesangial expansion compared to LF-fed mice, whereas finerenone treatment significantly reduced mesangial enlargement in WD-fed animals (Fig. 2A). Immunofluorescence staining for synaptopodin, a marker of podocyte integrity, revealed a notable decrease in WD-fed mice compared to LF-fed controls; finerenone intervention preserved synaptopodin expression and thereby mitigated podocyte damage (Fig. 2B).

**Figure 2:**
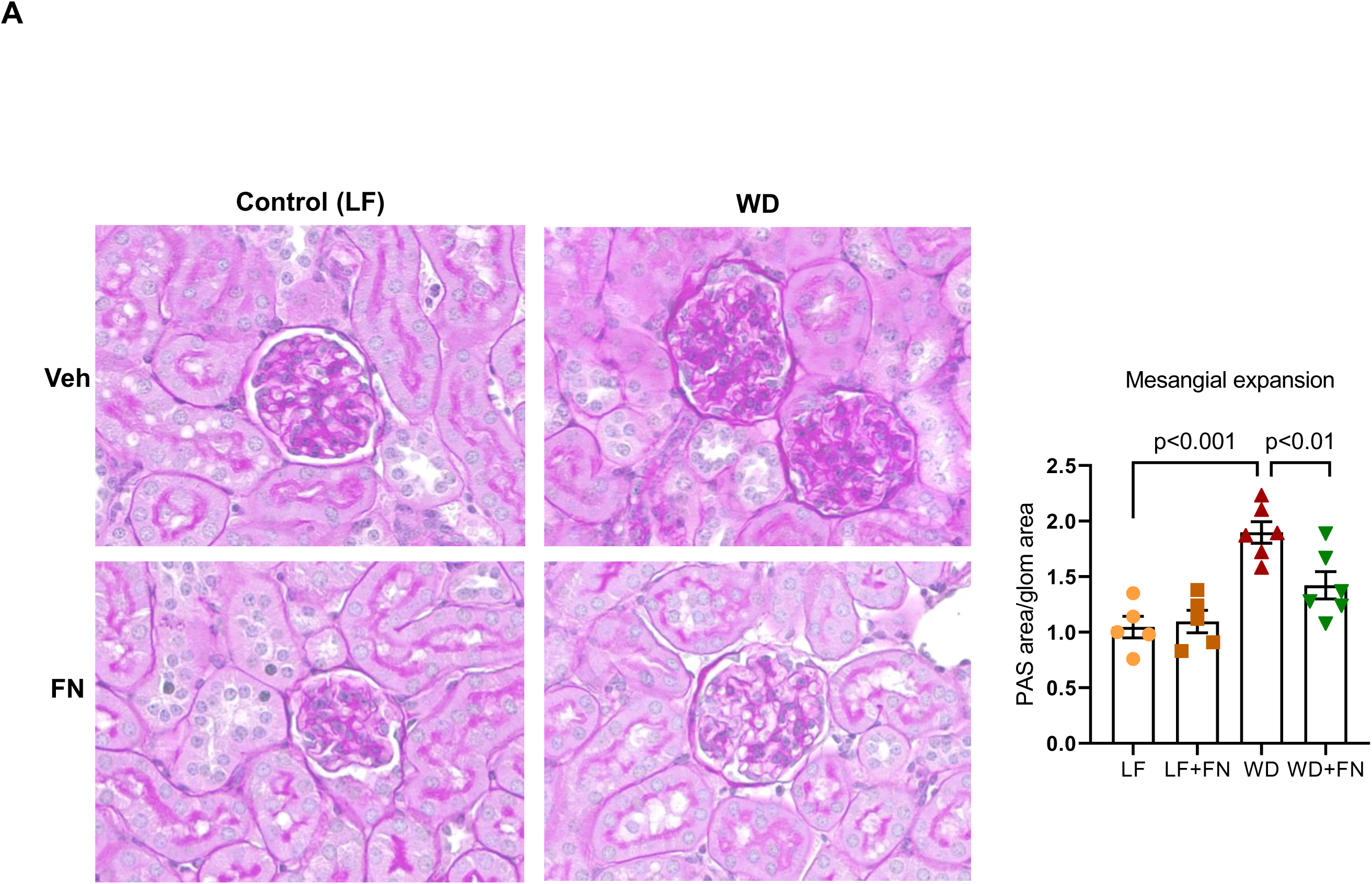
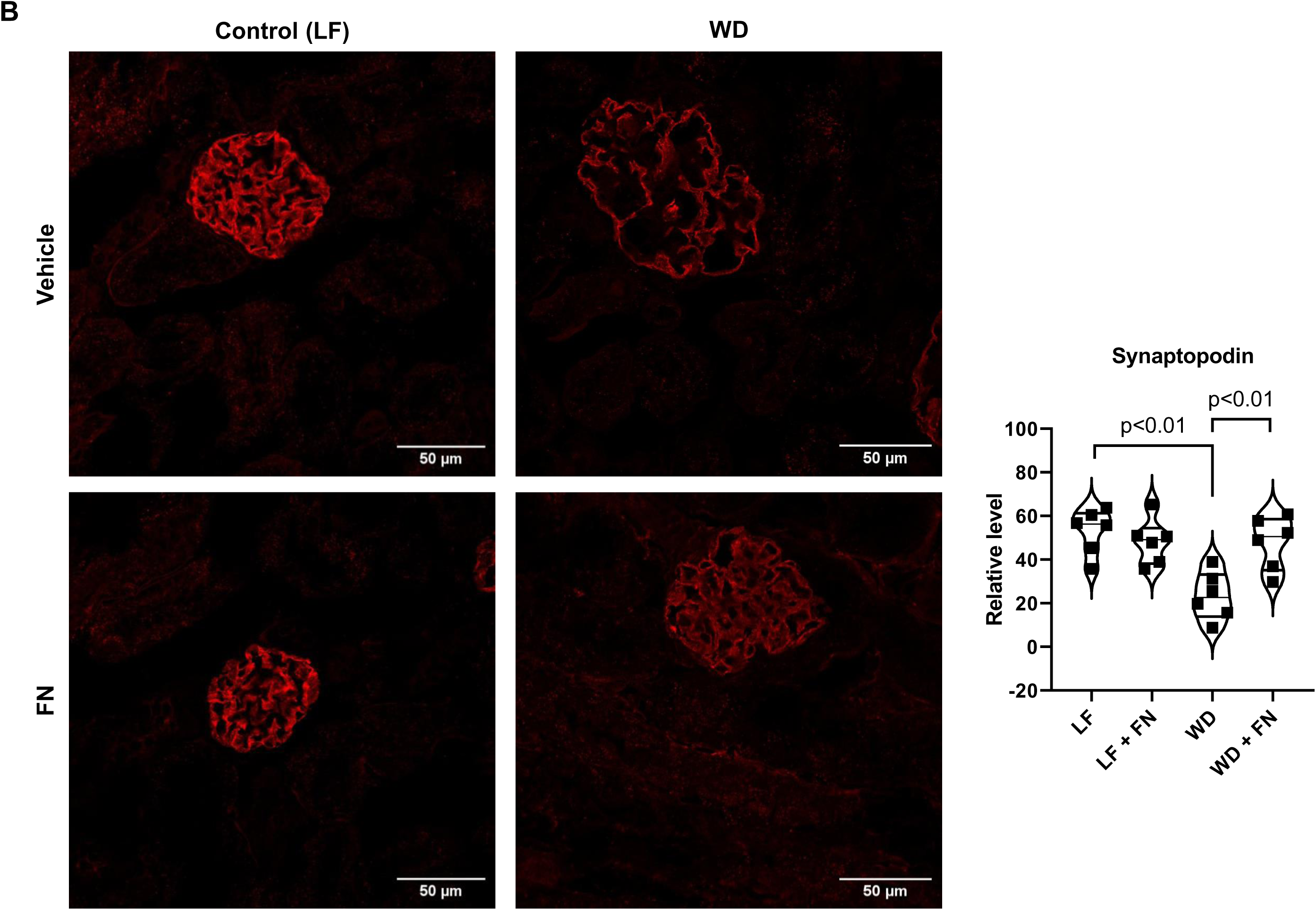
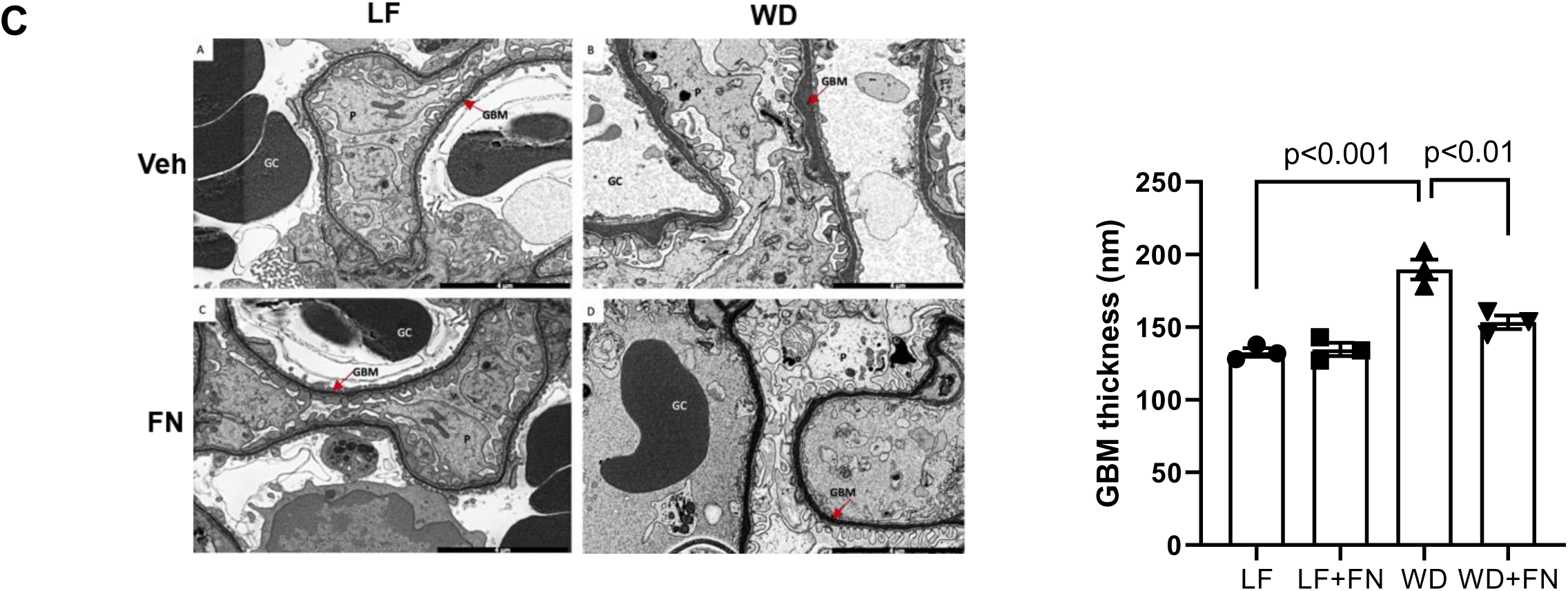
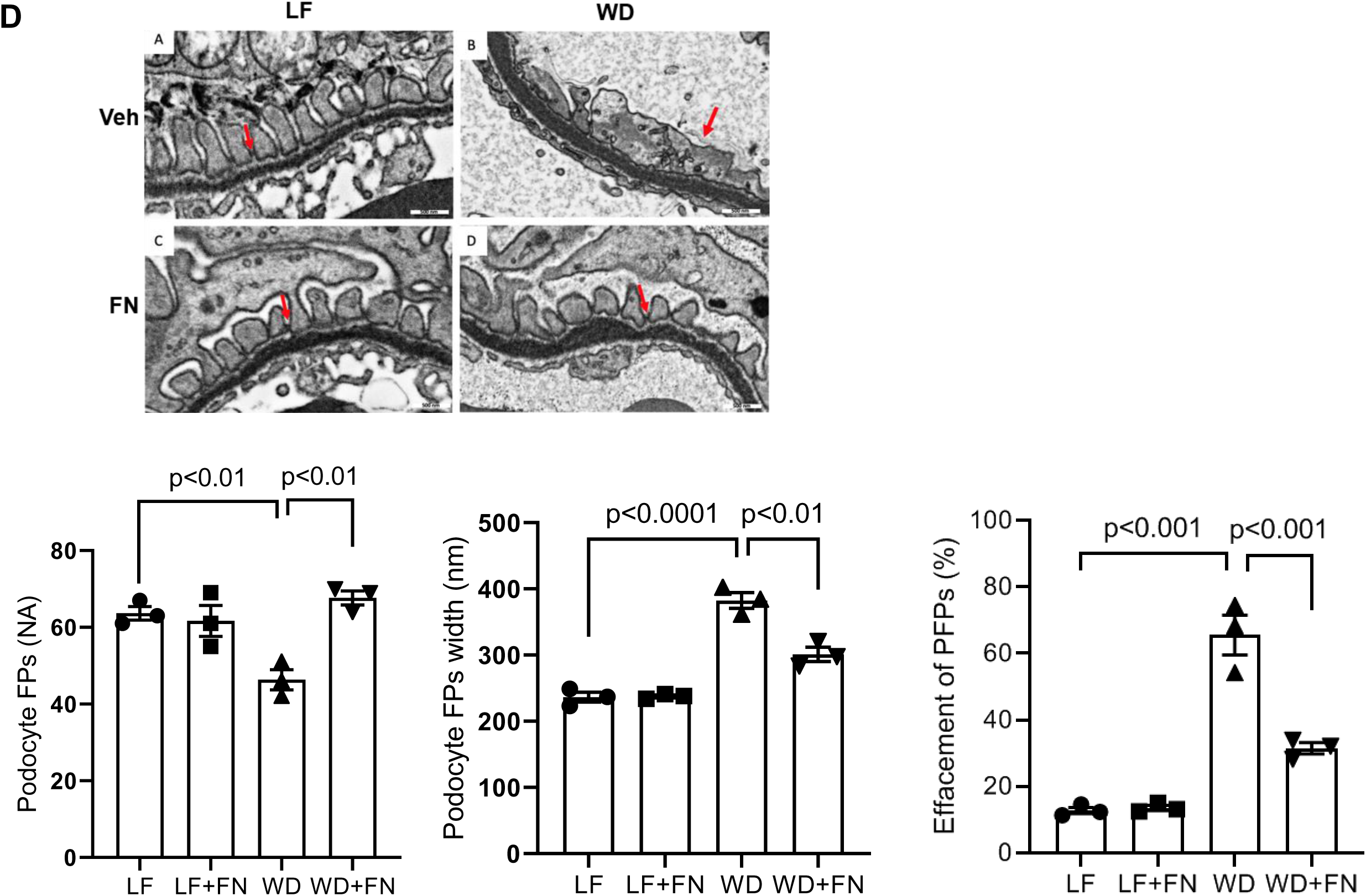
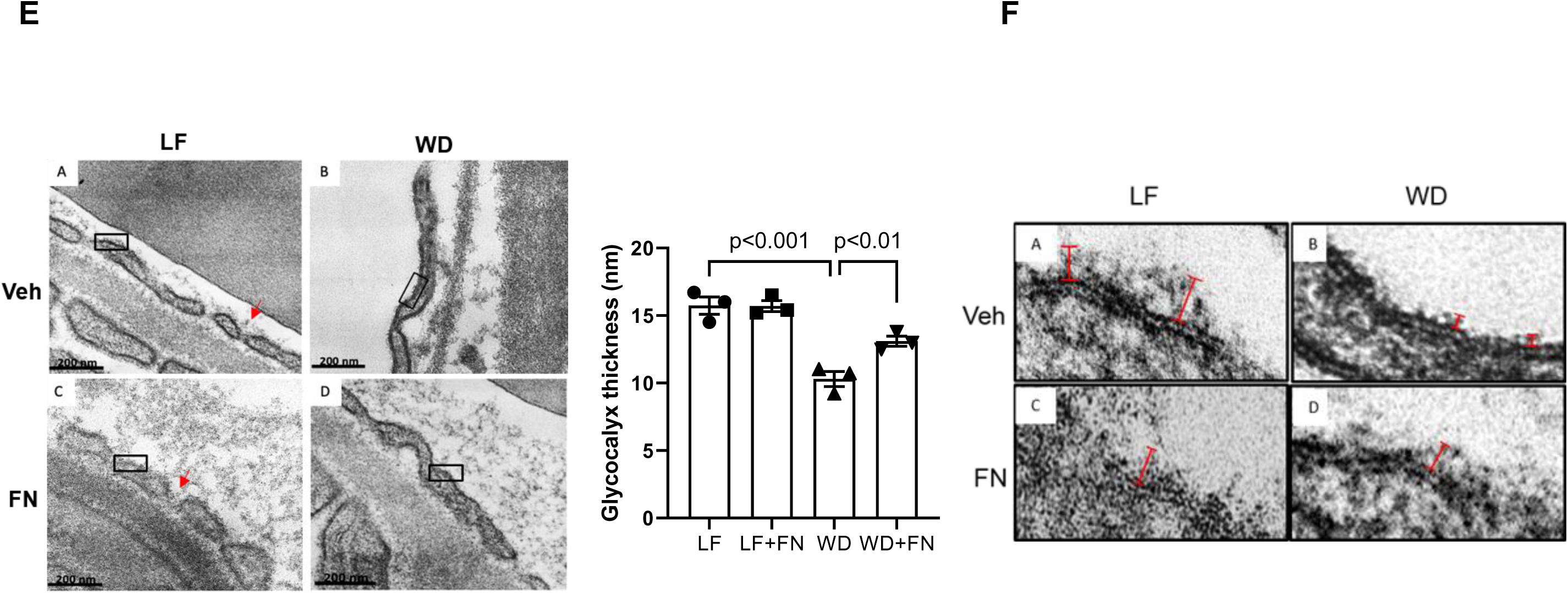
Finerenone treatment prevented glomerular histology, ultrastructure, and podocyte function in diet-induced obesity. A) Periodic Acid-Schiff (PAS) staining of kidney sections indicating the accumulation of extracellular matrix proteins within the glomerular mesangial area and positive staining per each glomerulus, n=6 per group B) Immunofluorescence staining of synaptopodin in the kidney and its relative levels representing the ratio of the synaptopodin positive area to the glomerular tuft area, n=6 per group. C) SEM images demonstrate the glomerular ultrastructure and glomerular basement membrane (GBM) thickness along with quantified bar graph. Magnifications x 35,000, n=3 per group. D) SEM images demonstrate the structure of podocyte foot processes (PFPs). Quantification performed as relative level of normal PFPs, width (nm) and assessed effacement of PFPs by identifying broadened, flattened, and fused processes with loss of slit diaphragms, n=3 per group E) TEM images illustrate the glycocalyx distribution in glomerular endothelial cells, measurements were taken from ten sites per capillary across five glomerular capillaries per animal. Magnifications x 80,000, n=3 per group F) Enlarged fragments within the highlighted rectangles further illustrate glycocalyx thickness (red line). The bar graph shows quantification of glycocalyx thickness (nm). n=3 per group. p-value indicated as p<0.01, p<0.001.

Electron microscopy was then employed to examine ultrastructural changes in the GBM and podocyte foot processes. Scanning electron microscopy (SEM) revealed GBM thickening in WD-fed mice, which was alleviated by finerenone supplementation (Fig. 2C). Moreover, WD-fed mice exhibited podocyte foot process effacement, widening, fusion, and loss of slit diaphragms, but these abnormalities were prevented by finerenone (Fig. 2D). Consistent with prior reports indicating that mineralocorticoid receptor inhibition safeguards the endothelial glycocalyx in diabetic conditions (60), transmission electron microscopy (TEM) revealed significantly reduced glycocalyx thickness in WD-fed mice, an effect that was reversed by finerenone (Fig. 2E). Higher magnification TEM images further demonstrated decreased glycocalyx density and distribution in WD-fed mice, alterations that were similarly restored by finerenone (Fig. 2F). Collectively, these results demonstrate that finerenone significantly ameliorates WD-induced kidney damage, preserving both functional and structural integrity in the glomerulus and tubules.

### Finerenone treatment ameliorated renal fibrosis in western diet-induced obese mice

The accumulation of fibrillary collagens within the renal tubular interstitium is a well-documented feature in mice fed a high-fat diet (19, 61, 62). Picrosirius red (PSR) staining, followed by polarized light imaging and subsequent quantification, revealed elevated fibrillary collagen deposition in the tubular interstitial space of WD-fed kidneys compared to LF-fed controls. This increase was attenuated by finerenone treatment (Fig. 3A–C). Immunofluorescence staining for fibronectin and collagen type IV both key extracellular matrix components produced by mesangial cells that contribute to glomerulosclerosis showed significantly higher accumulation in WD-fed mice compared to LF-fed mice. However, finerenone administration prevented these increases (Fig. 3D-3E). Consistent with these findings, both RT-qPCR and western blot analyses revealed upregulated fibronectin transcript and protein levels in WD-fed kidneys, which were suppressed by finerenone (Fig. 3F). Furthermore, transcript levels of collagen-4 alpha chain 2 (Col4a2) and several pro-fibrotic markers including transforming growth factor-β (TGFβ), plasminogen activator inhibitor-1 (Pai1), connective tissue growth factor (CTGF), and tissue inhibitor of metalloproteinase 1 (Timp1) were assessed. WD-fed mice exhibited significantly elevated levels of Col4a2, Pai1, TGFβ, CTGF, and Timp1 compared to LF-fed mice. Finerenone treatment lowered these indices to levels observed in LF-fed controls (Fig. 3G). These findings indicate that finerenone significantly attenuates kidney fibrotic remodeling in diet-induced obese mice, underscoring its therapeutic potential in preventing renal injury.

**Figure 3:**
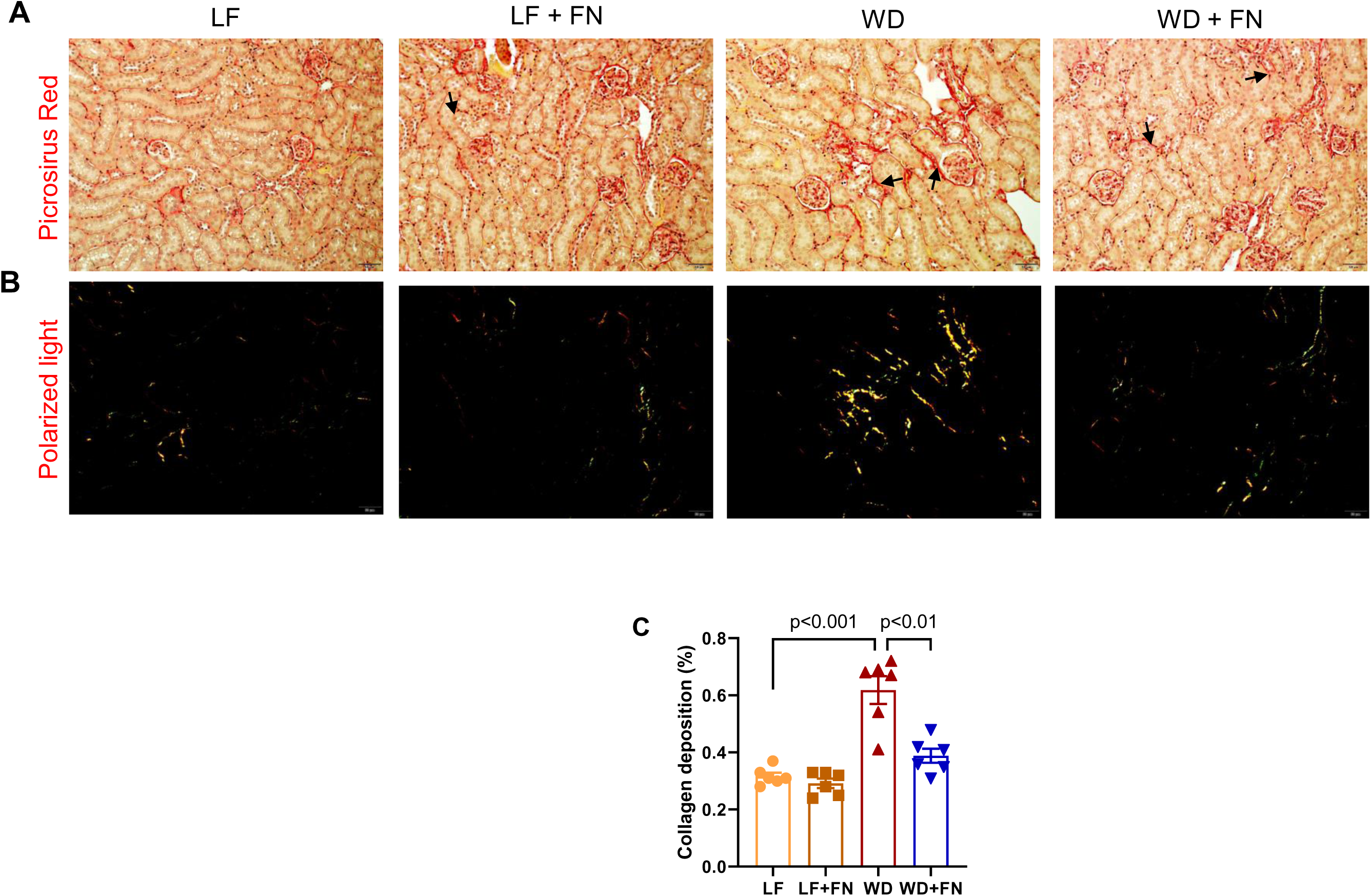
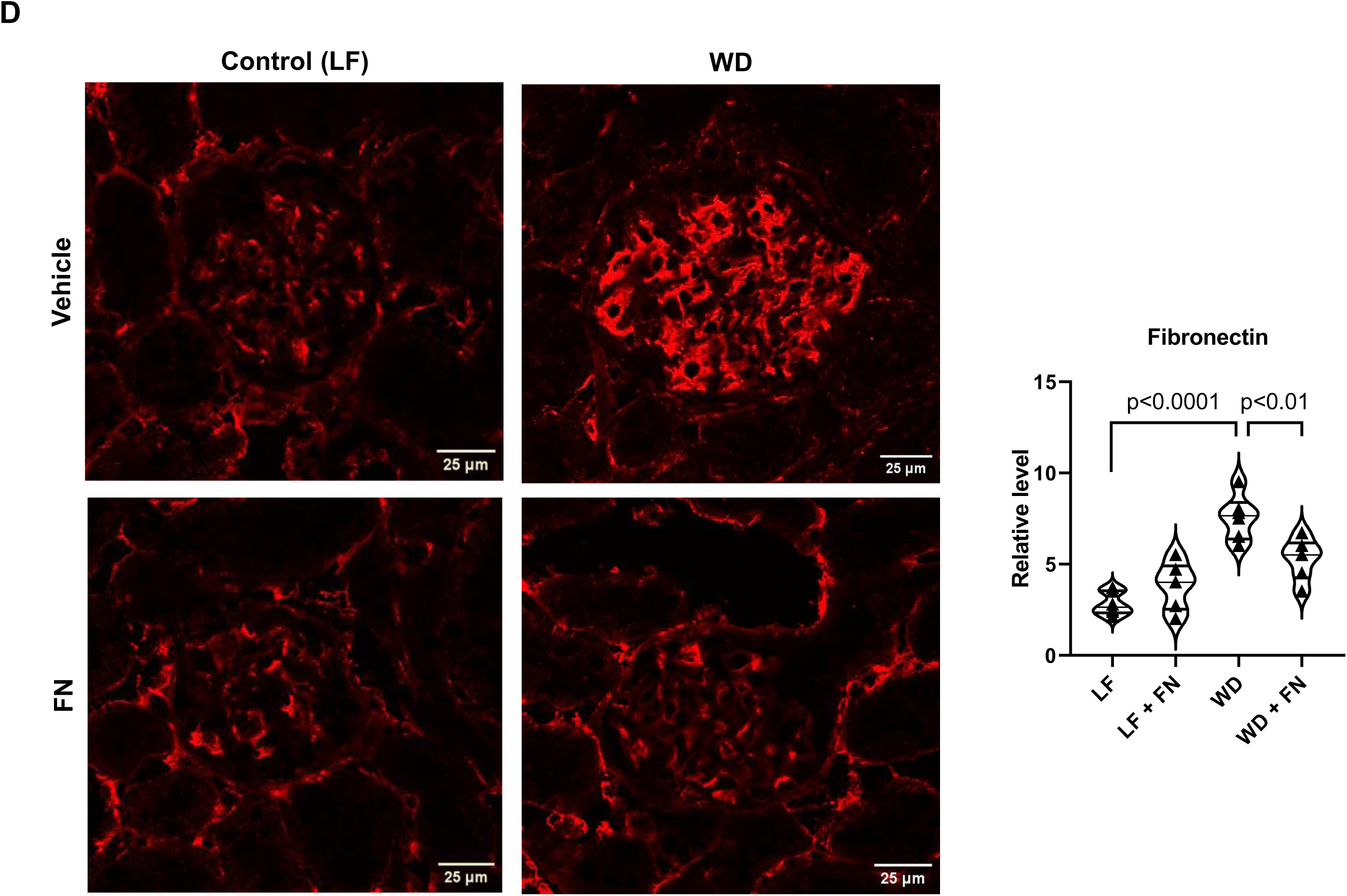
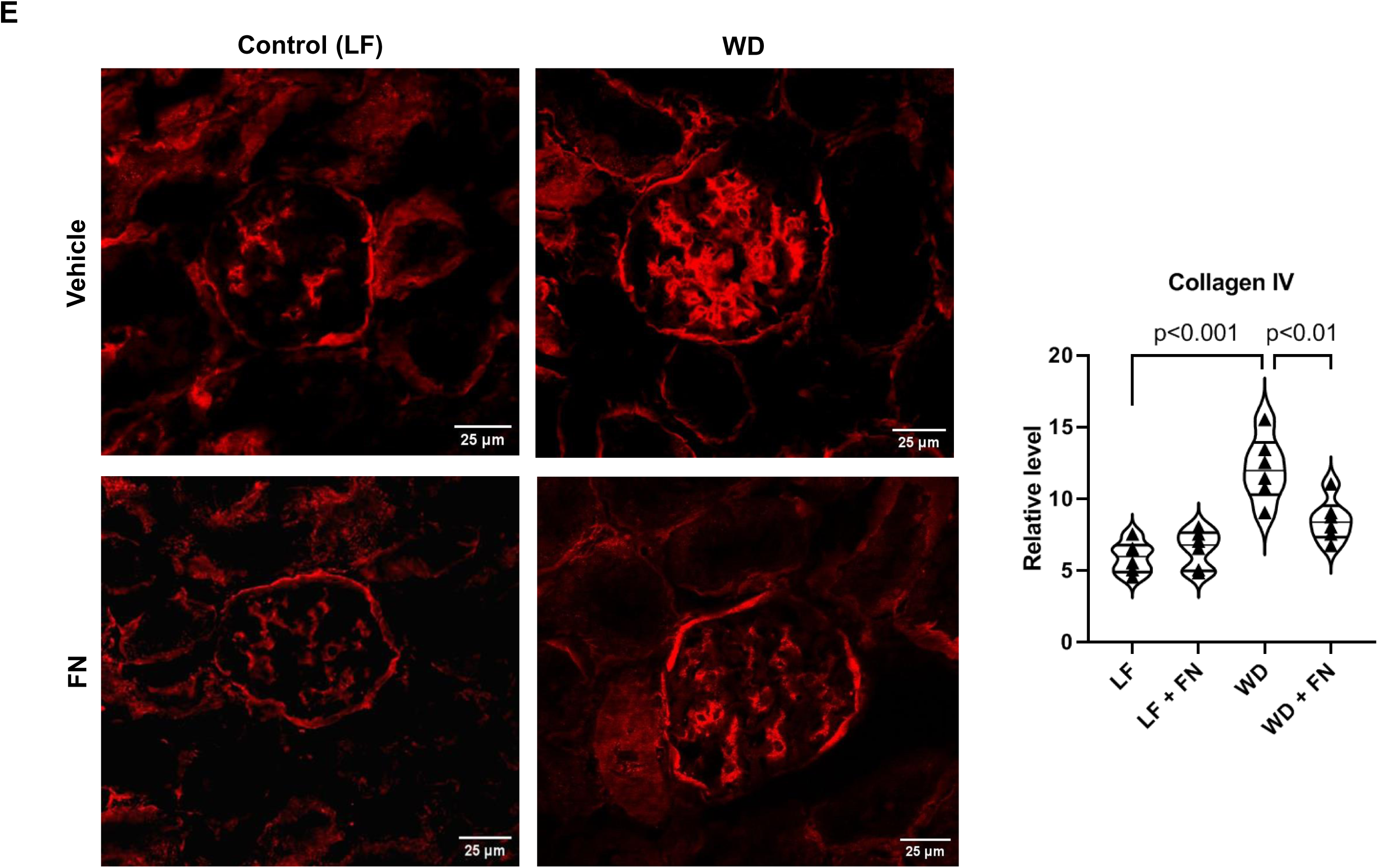
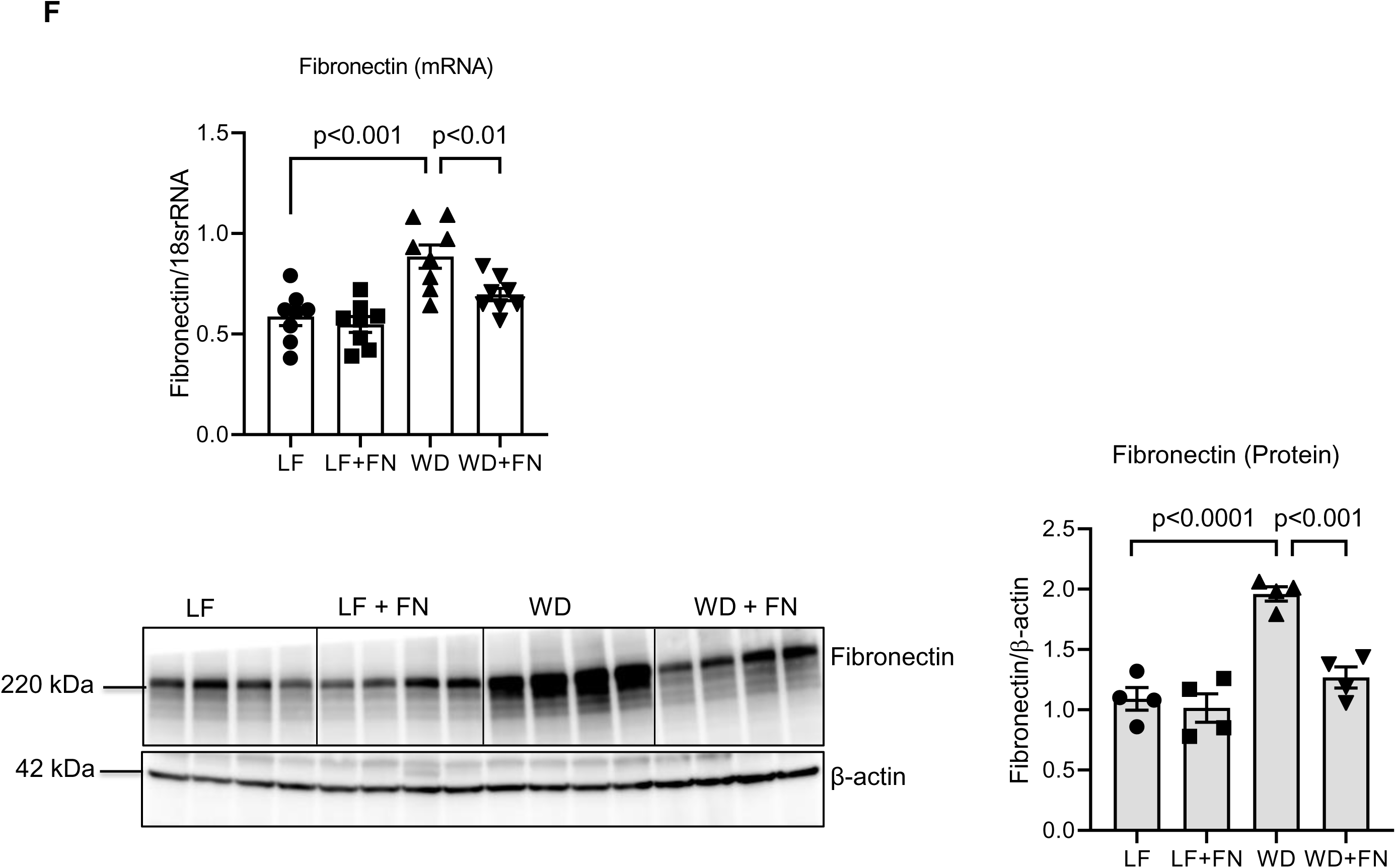
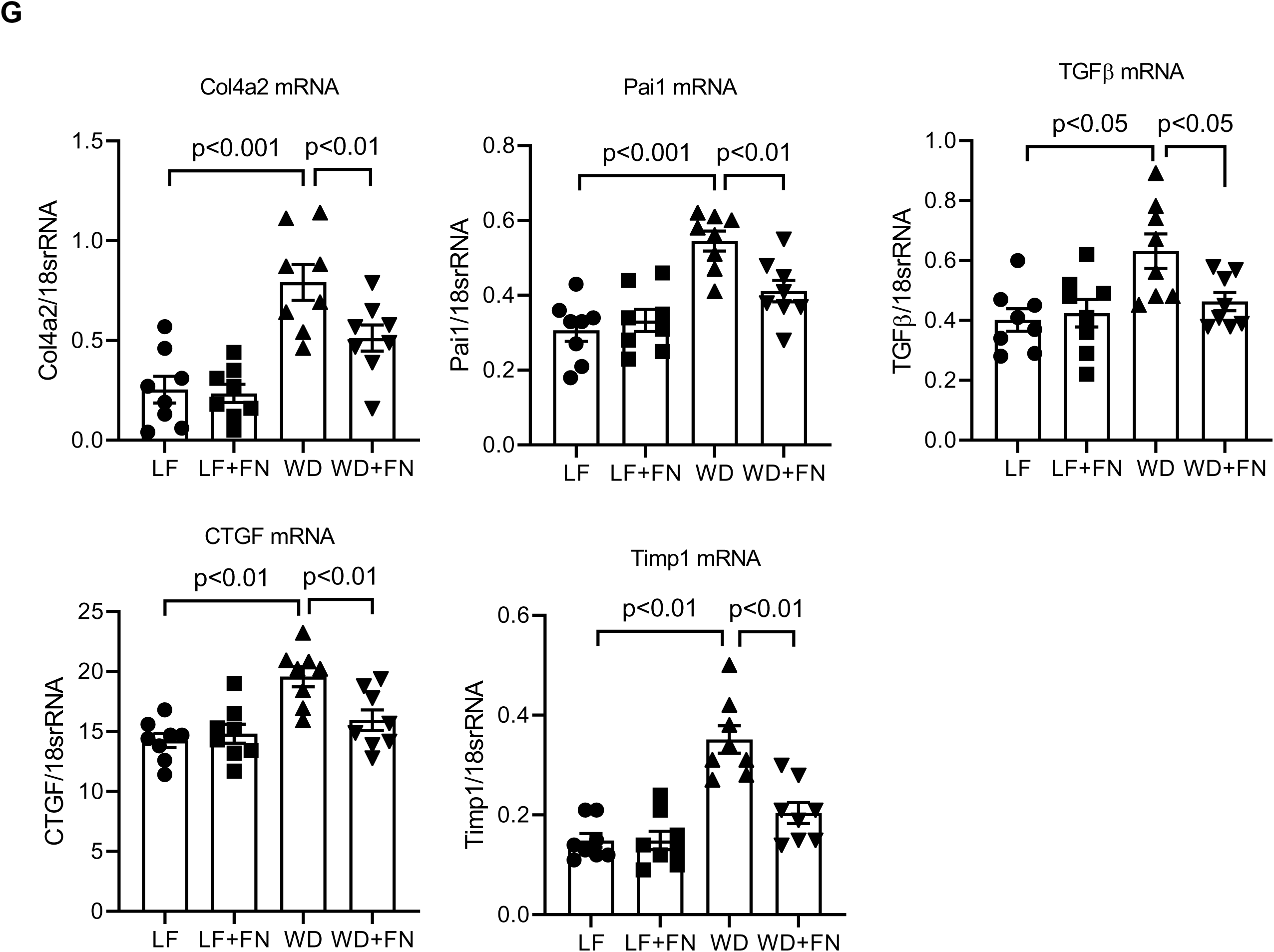
Finerenone treatment ameliorated Western diet-induced kidney fibrosis. A) Picrosirius red (PSR) staining highlights collagen deposition in the tubulointerstitial (arrow) and around glomeruli (arrowhead) n=6 per group. B) Representative images captured under polarized light microscopy from the same area illustrate collagen distribution. Scale Bar: 50 µm, n=6 per group C) Quantification of collagen deposition. Magnification x 20, n = 6 per group. Confocal microscopy images of kidney sections for D) fibronectin expression E) collagen-IV accumulation, and quantification of the positive staining area per each glomerular area. n=6 per group. F) qRT-PCR and immunoblot (every 2 samples in each group randomly pooled) analysis of fibronectin expression and quantification performed by normalizing to 18srRNA and β-actin respectively, n=8 per each group. G) qRT-PCR analysis of collagen-4 alpha chain2 (Col4a2), plasminogen activator inhibitor (PAI1), transforming growth factor-β (TGFβ), connective tissue growth factor (CTGF) and tissue inhibitor of metalloproteinases-1 (Timp1) and 18srRNA used as the housekeeping gene, n=8 per group. significance indicated as p-value (p<0.05).

### Finerenone decreases kidney triglyceride, cholesterol and ceramides

Previous studies have established that increased lipid accumulation in the kidney significantly contributes to the pathogenesis of kidney disease in mouse models of diet-induced obesity (61, 63, 64). To evaluate whether finerenone administration influences renal lipid accumulation, we quantified triglyceride and total cholesterol levels. Notably, WD-fed mice treated with finerenone showed a significant reduction in renal triglyceride levels. Total kidney cholesterol, which was markedly elevated in WD-fed mice compared to LF-fed controls, was also diminished by finerenone (Fig. 4A). In addition, we extracted lipids and analyzed ceramides by LC-MS. Levels of the long-chain ceramides C24:0 and C26:0 were elevated in kidneys from WD-fed mice compared to LF-fed mice but were significantly reduced following finerenone treatment (Fig. 4B). Our findings demonstrate that finerenone effectively reduces detrimental lipid accumulation in the kidneys of diet-induced obese mice, highlighting its potential to protect against obesity-related kidney damage

**Figure 4:**
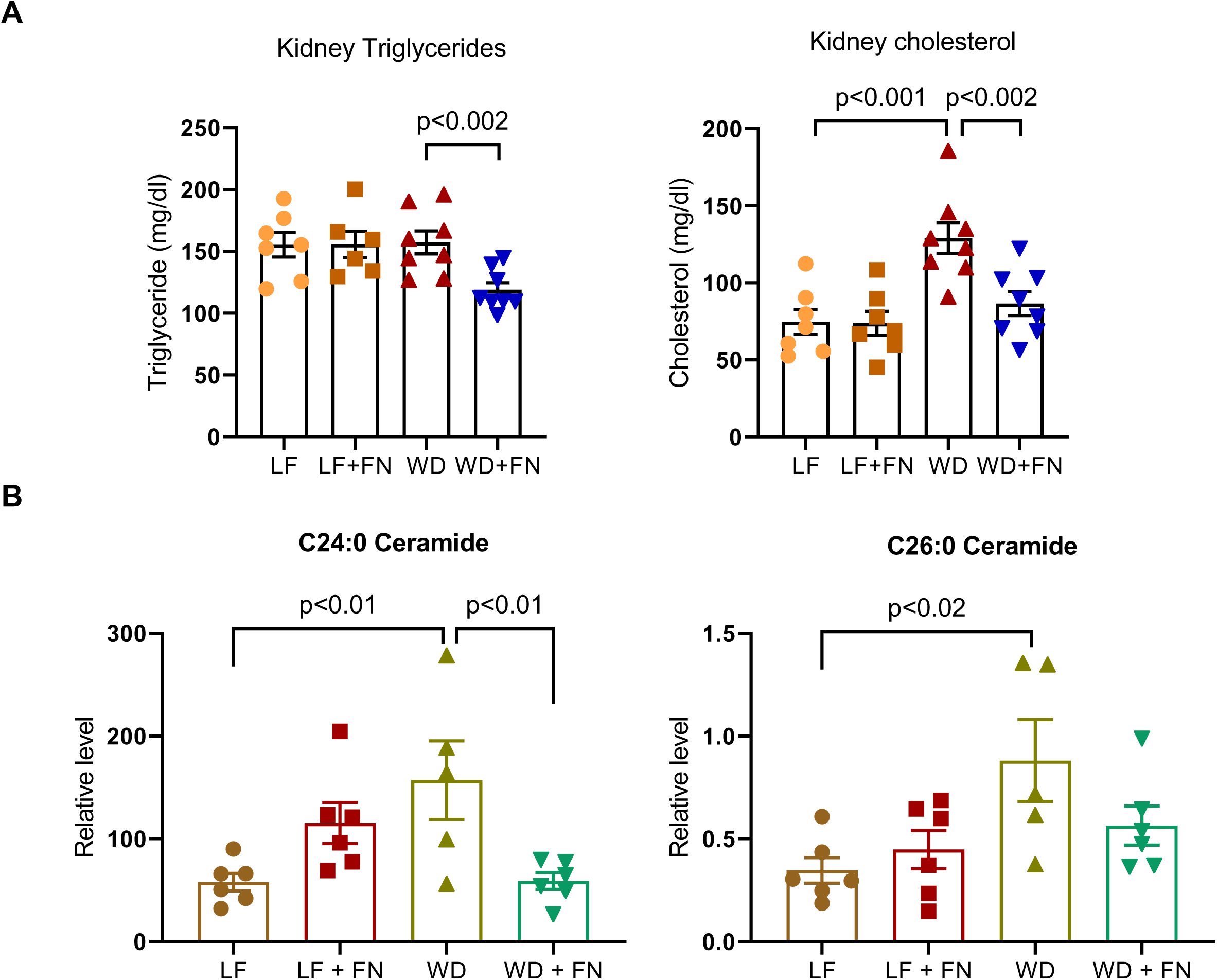
Finerenone treatment inhibited renal lipid accumulation in western diet-induced obesity. Total kidney A) triglycerides and cholesterol levels measured by colorimetric assay and bar graphs indicating the quantification, n = 6-8 mice per group. B) Tissue lipids were extracted, and concentrations were determined by liquid chromatography and mass spectrometry (LC-MS). Relative levels of long-chain ceramides C24:0 and C26:0, concentrations were quantified using peak-to-peak internal standards and ratios calculated with the tissue weight of individual samples, n = 5-6 per group. significance indicated as p<0.01 and p<0.02.

### Finerenone treatment attenuated the western diet-induced inflammation

To elucidate the role of inflammation in obesity-induced kidney disease, we examined the infiltration of CD45+ pan-leukocytes and CD68+ macrophages. Mice fed a western diet (WD) exhibited a significant increase in both CD45+ and CD68+ cells compared to low-fat (LF) controls. However, finerenone administration attenuated the accumulation of these inflammatory cells (Fig. 5A-B). Additionally, WD-fed mice showed a significant upregulation of inflammatory markers (MCP1, NLRP3 inflammasome) and DNA sensor pathways (cGAS, STING). Finerenone treatment reduced the mRNA abundance of these genes to near control levels (Fig. 5C). Consistent with transcript data, western blot analysis demonstrated elevated STING protein in WD-fed mice, which was suppressed by finerenone. STAT3 is a well-established inflammatory signaling mediator involved in the progression of obesity- and diabetes-related kidney disease (65, 66). Notably, WD feeding led to increased levels of phosphorylated STAT3 (p-STAT3), whereas finerenone significantly reduced p-STAT3 without affecting LF-fed controls (Fig. 5D). Taken together, finerenone inhibits obesity-driven inflammatory cell infiltration and proinflammatory signaling cascades in the kidney, revealing its potential to inhibit inflammation-associated renal damage in diet-induced obesity.

**Figure 5:**
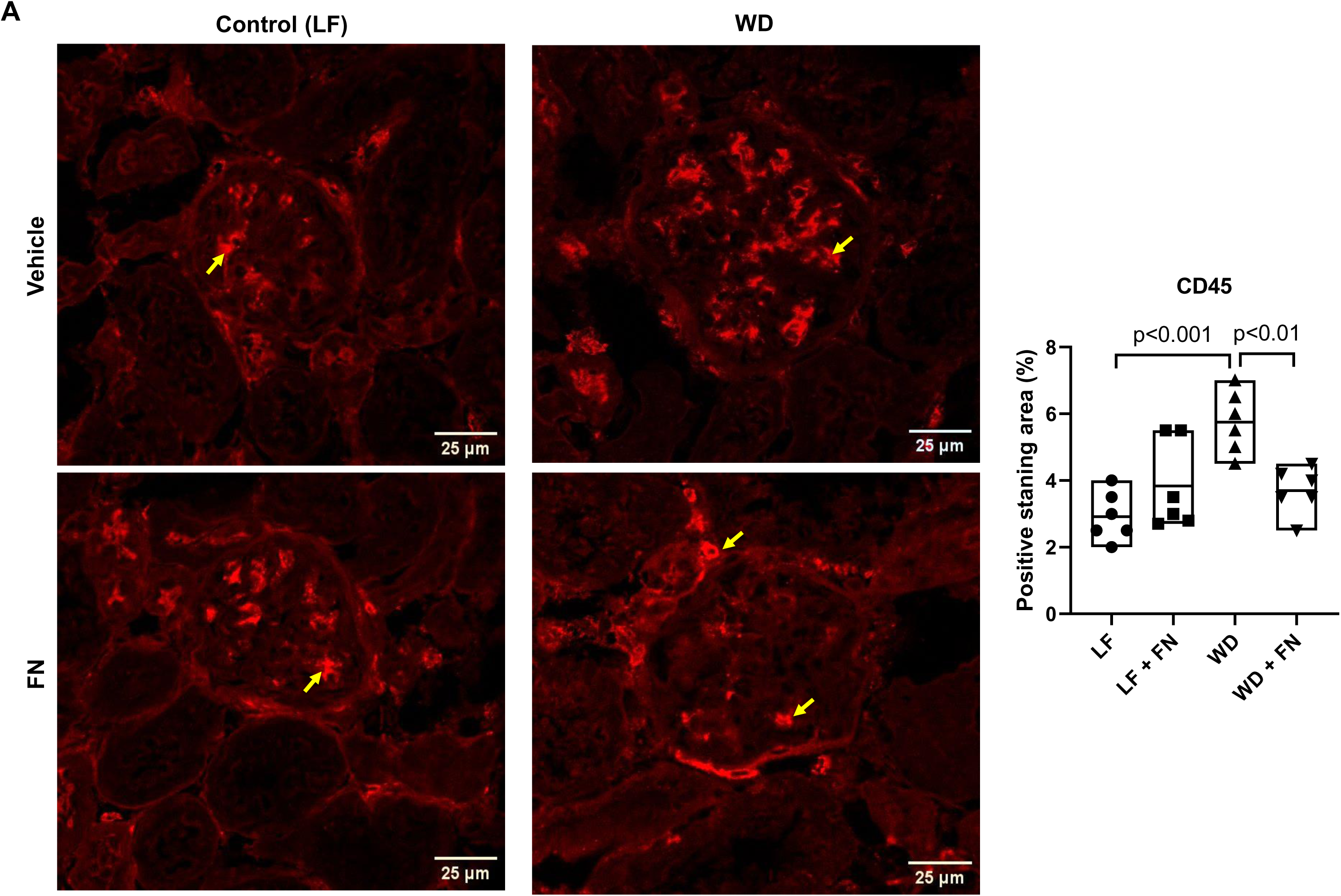
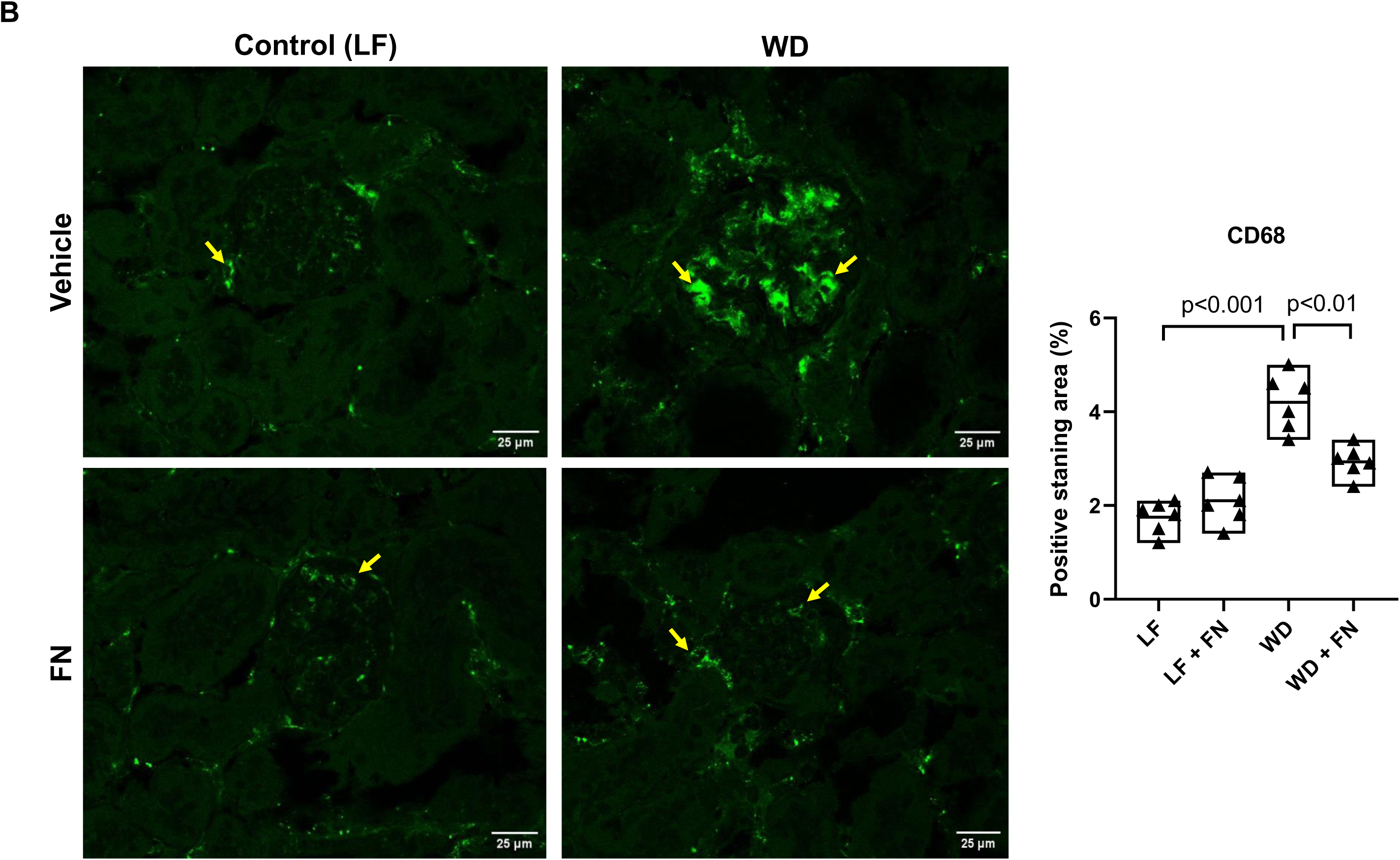
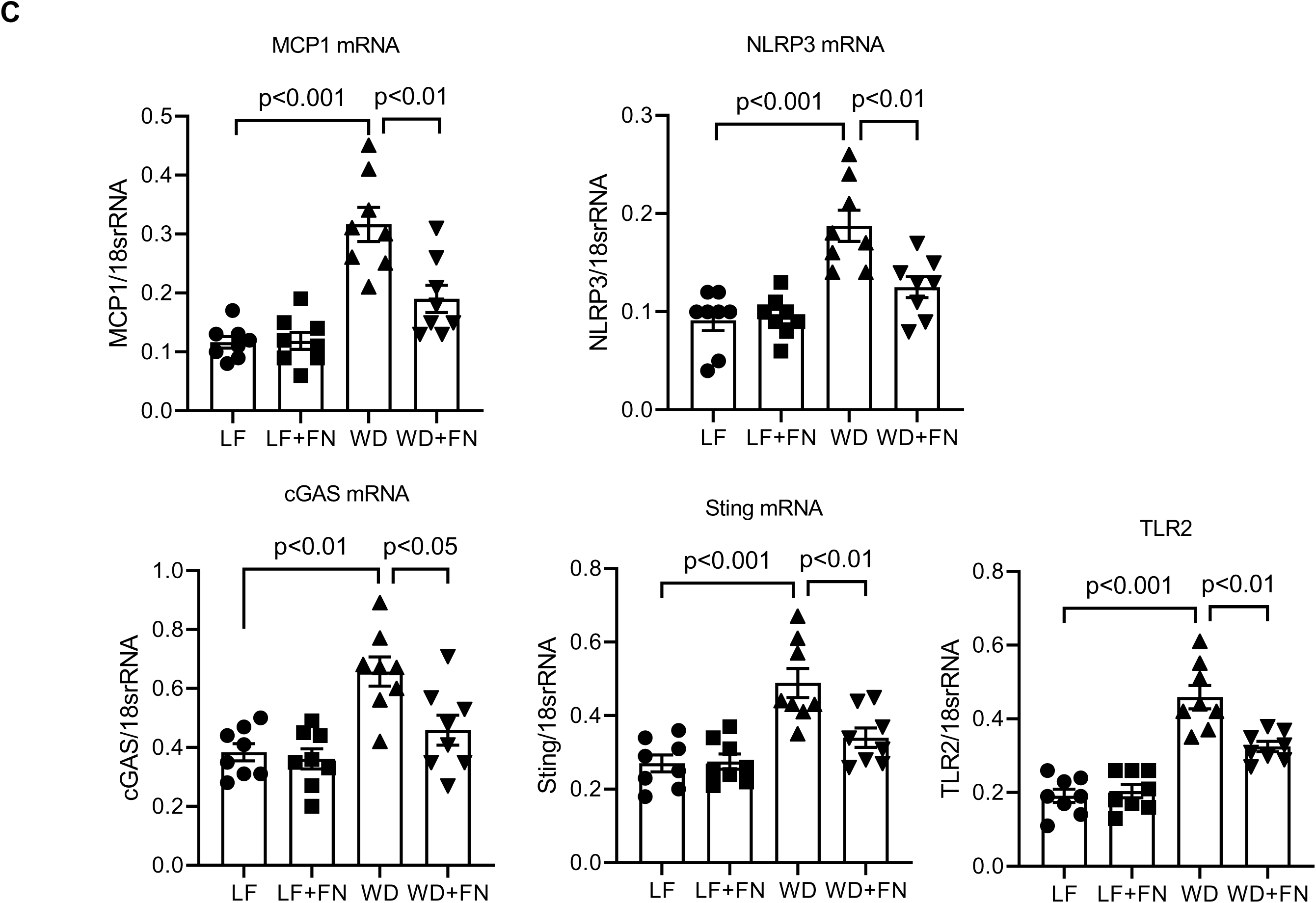
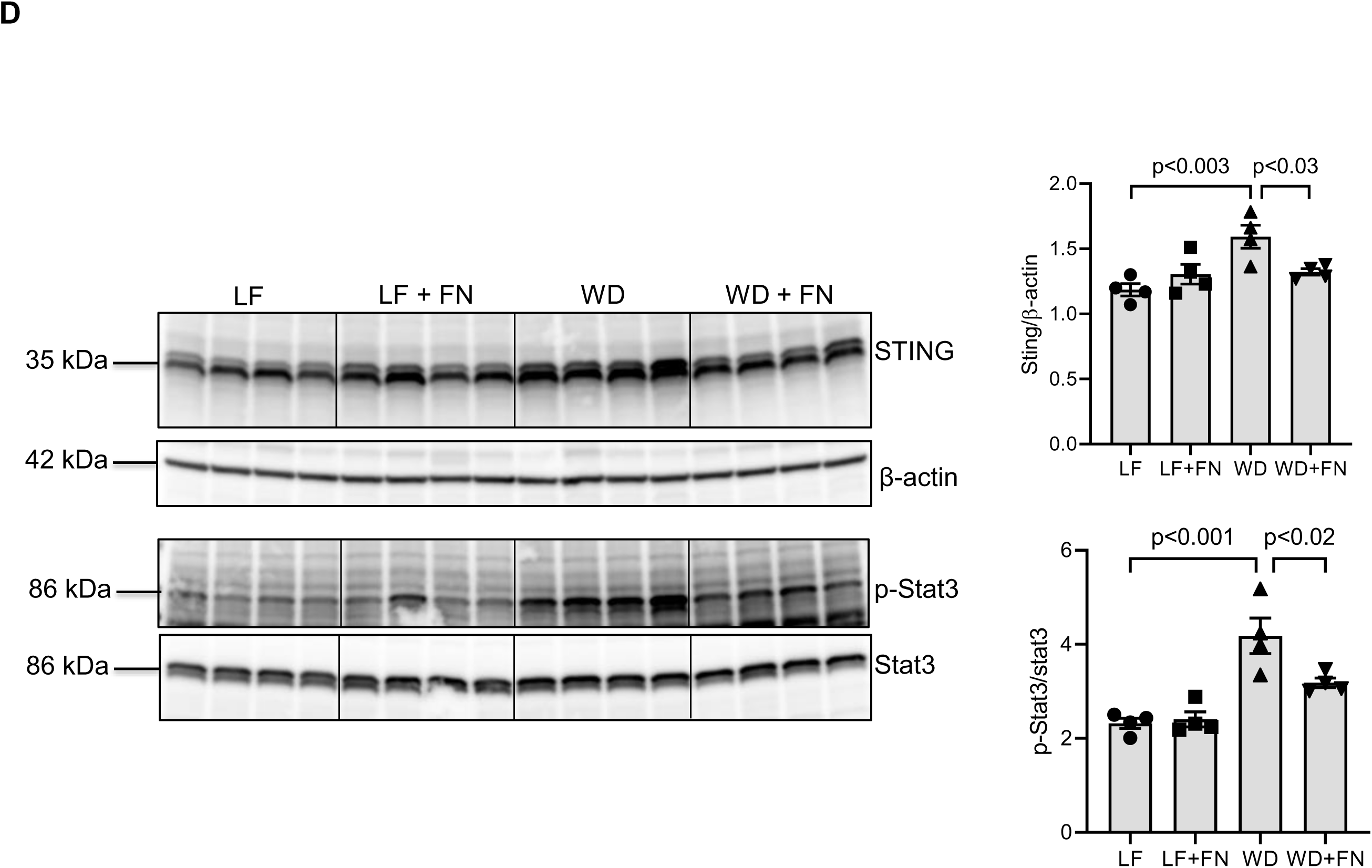
Finerenone administration reduces the inflammation in the kidneys of western diet-induced obesity. Immunofluorescence staining for A) CD45 as pan leukocyte marker, B) CD68 as macrophage marker in kidney sections, and percentage of positive staining (indicated as yellow arrow) area quantified, n=6 per group (A and B). C) qRT-PCR showing the gene expression of pro-inflammatory cytokine MCP1 and NLRP3 inflammasome and nucleotide sensors cGAS, sting, and TLR2, n=8 per group. D) immunoblot analysis of sting and phosphorylated stat3 and total stat3. n=8 per group (every 2 samples in each group randomly pooled).

### Finerenone ameliorated the mitochondrial morphology and function

Diet-induced obesity leads to mitochondrial dysfunction, characterized by impaired energy metabolism, altered mitochondrial morphology, and reduced oxidative phosphorylation (OXPHOS), all contributing to kidney disease (67–69). To investigate these changes, we used label-free fluorescence lifetime imaging microscopy (FLIM) to measure the ratio of free to bound NADH, thereby estimating the relative rates of glycolysis versus OXPHOS (70–73). A lower free/bound NADH ratio indicates reduced glycolysis and enhanced OXPHOS, while a higher ratio reflects increased glycolysis and diminished OXPHOS. Specifically, we calculated the free-to-bound NADH ratio using the distinct autofluorescence decay rates of NADH in its free or protein-bound forms. We observed a significant increase in the free/bound NADH ratio in kidney sections from WD-fed mice compared to LF-fed controls. However, this ratio decreased in WD-fed mice receiving finerenone and remained unchanged in LF-fed mice treated with finerenone (Fig. 6A).

**Figure 6:**
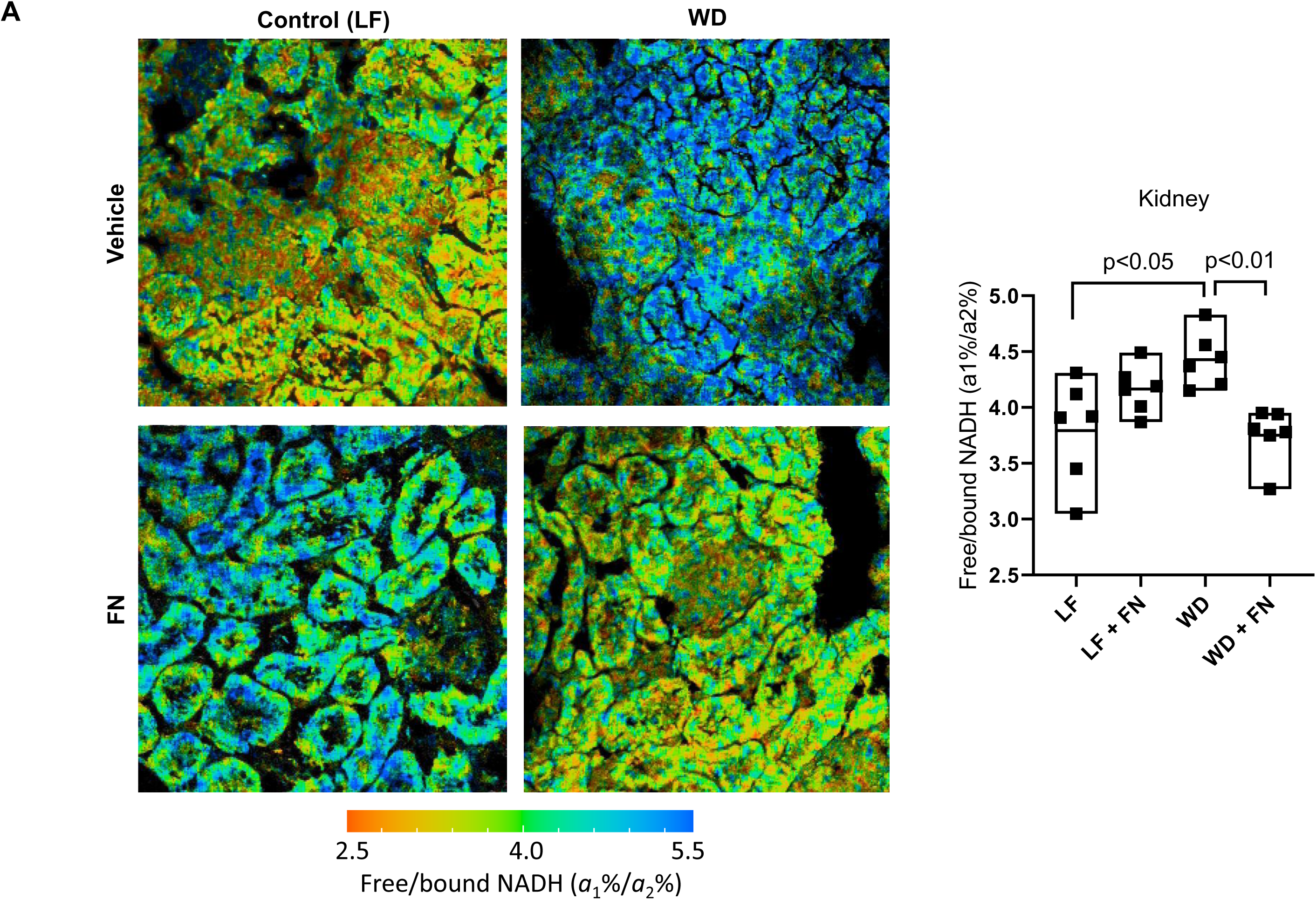
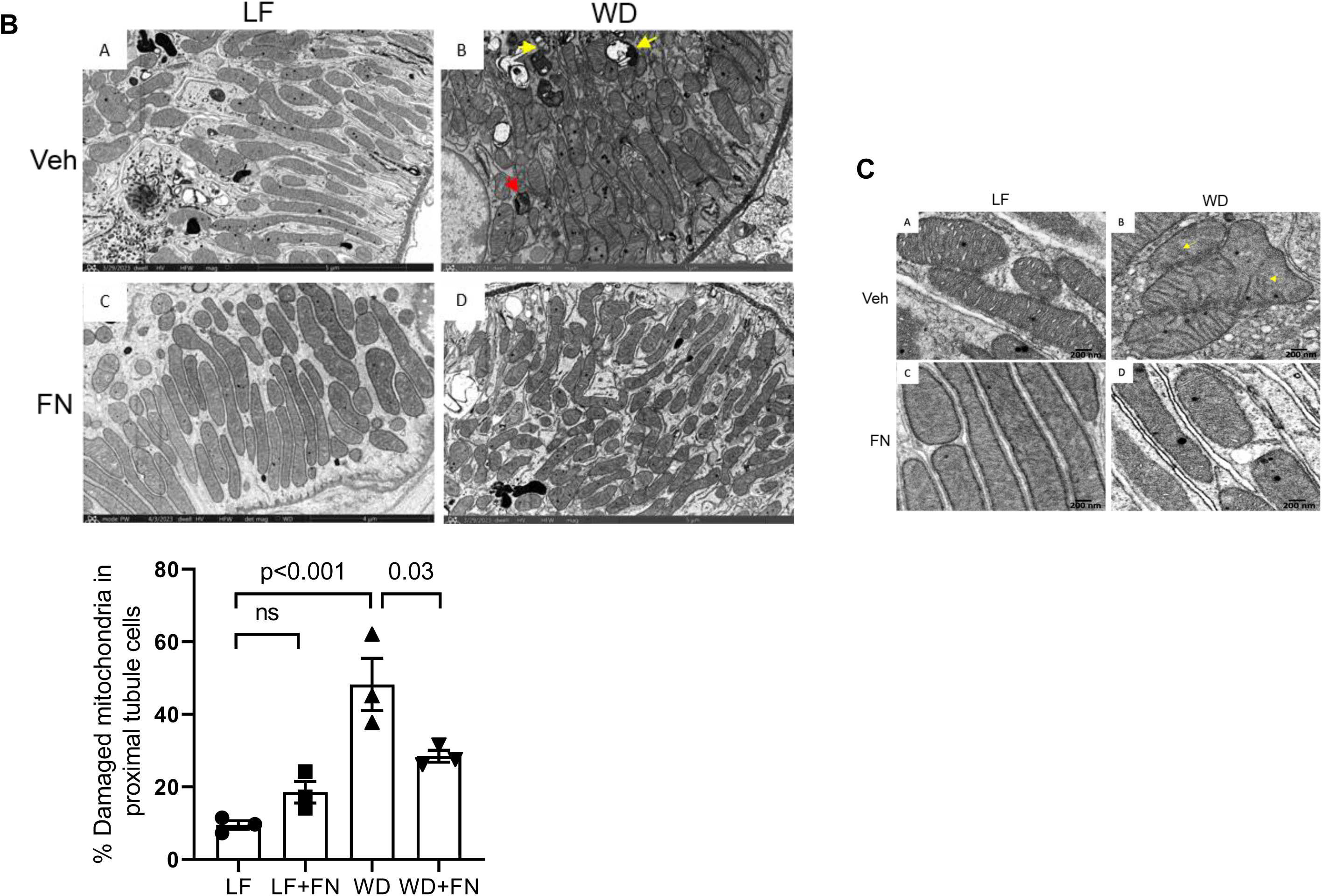
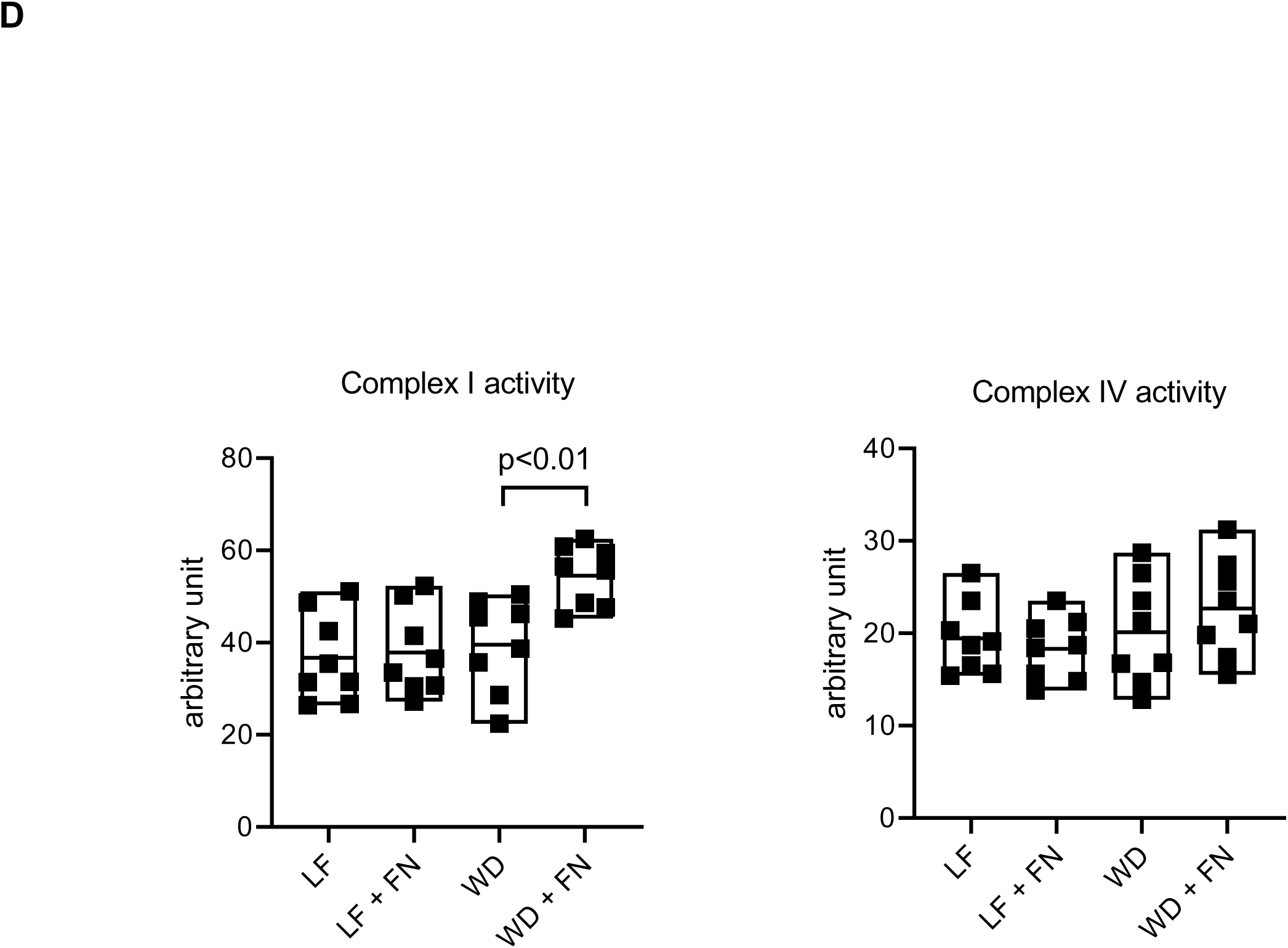
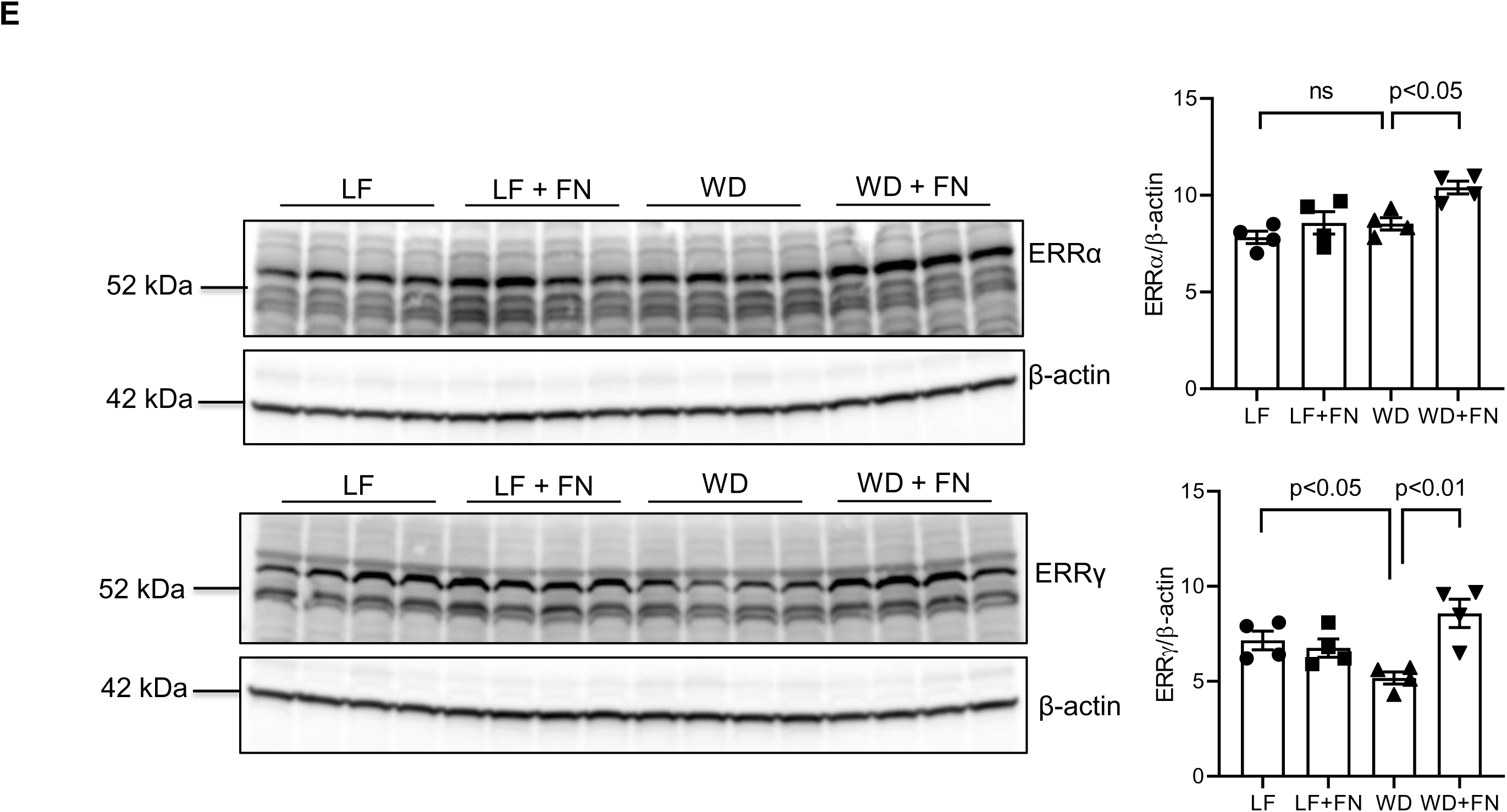
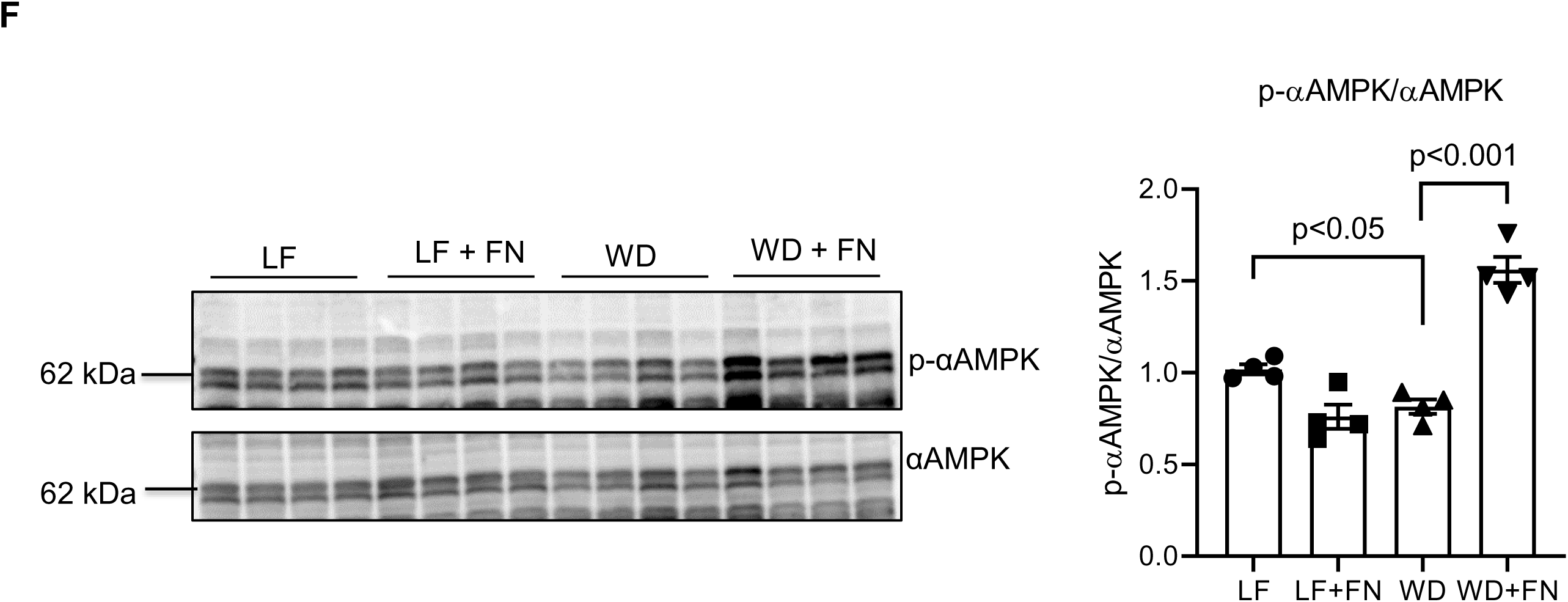
Finerenone treatment improved mitochondrial oxidative phosphorylation and morphology. A) Label-free two-photon imaging microscopy was performed to determine the NADH fluorescence lifetime and measured free versus bound NADH ratios to indicate the glycolysis versus oxidative phosphorylation (OXPHOS). NADH fluorescence lifetime was quantified using the phasor approach, n=6 per group. B) SEM images illustrate mitochondria in proximal tubule epithelial cells across different groups: low-fat diet (LF, panel A), western diet (WD, panel B), and finerenone treated mice under LF (panel C) and WD (panel D) conditions. Magnification: ×35,000. C) Images acquired under higher magnification shows mitochondria ultrastructure. Percentage of damaged mitochondria quantified in proximal tubules and indicated as bar graph. Magnification x 50,000, n=3 per group. Total kidney was homogenized and measured for D) Total complex I and complex IV activity using standard microwell plate assay, n=8 per group. Immunoblot images showing the protein expression of E) nuclear receptors ERRα and ERRγ, and F) phosphorylated αAMPK and total αAMPK n=8 per group (every 2 samples in each group randomly pooled).

Consistent with these findings, electron microscopy revealed that WD-fed mice exhibited significant mitochondrial disorganization, with chaotically distributed and disoriented mitochondria, in contrast to the well-organized mitochondrial arrangement in LF-fed mice. Structural abnormalities in WD-fed mice included cristae fragmentation, homogenization, and swelling of the mitochondrial matrix, along with an increased presence of secondary lysosomes closely associated with damaged mitochondria (yellow arrows) (Fig. 6B, 6C). Finerenone treatment in WD-fed mice effectively improved mitochondrial organization, enhancing both distribution and cristae architecture restoration. In LF-fed mice treated with finerenone, mitochondria remained well-organized and structurally intact, indicating that finerenone had no adverse effects on normal mitochondrial morphology (Fig. 6B). Higher-magnification imaging provided deeper insights into mitochondrial ultrastructure (Fig. 6C). WD-fed mice exhibited cristae disorganization, loss of tight packing, and matrix swelling, compared to the densely packed cristae with regular spacing observed in LF-fed mice (Fig. 6C, Panel A and B). In WD-fed mice treated with finerenone, mitochondrial integrity was largely restored, showing more organized cristae, reduced swelling, and overall structural improvement (Fig. 6C Panel D). Morphometric analysis revealed a significant reduction in the number of damaged mitochondria (Fig. 6C).

Conversely, finerenone administration had no effect on mitochondrial morphology in LF-fed mice, as their mitochondria remained structurally normal (Fig. 6C, Panel C), nor did it affect the activity of mitochondrial complexes I or IV (Fig. 6D).

Since finerenone enhanced mitochondrial OXPHOS, we next explored possible molecular mechanisms. Previous studies have identified estrogen-related receptors α and γ (ERRα and ERRγ) as critical regulators of mitochondrial OXPHOS and fatty acid oxidation (74, 75). Western blot analysis revealed that the nuclear receptor ERRγ was downregulated in WD-fed mice, with no change in ERRα. Notably, finerenone treatment restored both ERRα and ERRγ expression levels in WD-fed mice but did not alter expression in LF-fed mice (Fig. 6E). Given the recognized role of AMPK in renal cell dysfunction under diet-induced obesity (63), we further examined whether mineralocorticoid receptor inhibition via finerenone activates AMPK signaling. WD-fed mice showed diminished phosphorylated AMPKα (p-AMPKα) relative to total AMPKα, which was significantly restored by finerenone. In contrast, p-AMPKα levels in LF-fed mice were unaffected by finerenone treatment (Fig. 6F). These findings underscore that finerenone effectively counteracts diet-induced mitochondrial dysfunction in the kidneys by enhancing OXPHOS, preserving mitochondrial integrity, and restoring key regulatory pathways (ERR and AMPK), thereby mitigating obesity-related renal damage.

Together, these findings demonstrate that finerenone treatment mitigated WD-induced mitochondrial structural damage and dysfunction.

## DISCUSSION

The present study demonstrates that finerenone provides renoprotection in a mouse model of western diet-induced obesity and insulin resistance by modulating mitochondrial homeostasis, inflammation, lipid metabolism and fibrosis. Western diet-fed mice exhibited significant renal dysfunction, marked by increased albuminuria, kidney injury molecule-1 (KIM1), and NGAL and structural abnormalities, including mesangial matrix expansion, podocyte injury, and renal fibrosis, all of which were mitigated by finerenone treatment.

Our data is consistent with previous study in a mouse model of high salt fed diabetic db/db-UNX induced hypertension, showing that finerenone treatment improved albuminuria and glomerular lesions (76). Hypertensive db/db-UNX mouse model shows more robust kidney damage and has clinical relevance to hypertensive diabetic patients with progressive kidney damage leading to kidney failure (76). However, our model is highly relevant to obesity patient population usually with early stage kidney lesions. Finerenone also reduced pro-inflammatory cytokines, innate immunity pathway activation, and fibrosis markers, lowered kidney cholesterol levels, and restored mitochondrial oxidative phosphorylation. Finerenone uniquely enhanced nuclear receptor ERRγ expression and corrected mitochondrial and ultrastructural abnormalities in WD-fed kidneys. These findings highlight finerenone’s multifaceted protective effects on inflammation, fibrosis, and mitochondrial health in WD-induced kidney injury. Finerenone may hold promise as a therapeutic agent for western diet-induced obesity and insulin resistance related kidney damage.

The beneficial effects of finerenone are linked to the upregulation or stabilization of estrogen-related receptor gamma (ERRγ) by modulating mitochondrial biogenesis and oxidative metabolism in renal cells. Enhanced ERRγ signaling improves mitochondrial integrity and function, attenuating lipid accumulation and reducing renal lipotoxicity. Concurrently, these metabolic improvements diminish proinflammatory signaling cascades characterized by reduced levels of proinflammatory cytokines and less infiltration of immune cells thereby diminishing renal inflammation and the progression of kidney injury. MR antagonism effectively prevents glomerular endothelial glycocalyx (GEnGlx) damage, reduces matrix metalloproteinase (MMP) activity, and restores glomerular albumin permeability in early diabetic nephropathy (DN). that supports MR antagonism as a promising therapeutic strategy to reduce renal damage (60).

In this study, finerenone treatment was initiated after 14 weeks of exposure to a western diet (WD), during which mice develop albuminuria as well as kidney and vascular damage. This intervention represents a treatment strategy to assess the efficacy of finerenone in mitigating kidney damage progression in a diet-induced obesity mouse model. Finerenone effectively suppressed the upregulation of key kidney damage markers, including Kim-1, NGAL, TBARS, and albuminuria. Furthermore, it conferred protection against podocyte injury by regulating the actin cytoskeleton protein synaptopodin. Obesity-induced damage to glomerular podocytes has been identified as a primary driver of obesity-related kidney disease (61). High-fat diet (HFD) administration in C57BL/6J mice, prone to diet-induced obesity, increases renal SREBP-1 and SREBP-2 activity and their target enzymes, leading to lipid accumulation, glomerulosclerosis, and proteinuria (61). The protective role of the SGLT2 inhibitor dapagliflozin in reducing kidney and liver complications associated with obesity in western diet-fed mice also supports our findings (64). High fat diet induced kidney disease is marked by the accumulation of lipid vacuoles predominantly in proximal tubular cells, accompanied by impaired mitochondrial morphology, lysosomal dysfunction and disrupted autophagy, suggest that a western diet induces multifaceted renal complications through distinct cellular and metabolic pathways (63, 64). Notably, finerenone not only ameliorated metabolic dysregulation but also prevented diet-induced podocyte injury, further underscoring its renoprotective effects in obesity-associated kidney disease.

The western diet (WD) exacerbates kidney injury, inflammation, and fibrosis through mechanisms that are not entirely attributable to its effects on glucose metabolism. Studies in patients with kidney disease have identified key pro-fibrotic biomarkers, including transforming growth factor-β (TGF-β), monocyte chemoattractant protein-1 (MCP-1), and matrix metalloproteinase-2 (MMP-2), which are strongly associated with fibrosis development and correlate with worsening renal function (77). Preclinical models have been instrumental in elucidating the role of the mineralocorticoid receptor (MR) in the progression of fibrosis and CKD, as well as evaluating the efficacy of finerenone in mitigating renal fibrosis. In the DOCA-salt rat model of CKD, finerenone significantly reduced the renal mRNA expression of the pro-fibrotic marker plasminogen activator inhibitor-1 (PAI-1) and alleviated renal fibrosis, as demonstrated by histopathological analysis (78–80). Additionally, finerenone has been shown to reduce the expression of other critical pro-fibrotic markers, including fibronectin, collagen IV, connective tissue growth factor (CTGF), TGF-β, and MCP-1 (36, 81).

Our findings align with previous studies, reinforcing the evidence that finerenone effectively reduces renal fibrosis and inflammation by targeting multiple pathways associated with these processes (80). A recent study identified key cell types and gene networks, including secreted phosphoprotein 1 (SPP1), interleukin-34 (IL-34), and platelet-derived growth factor subunit b (PDGFb), as drivers of hypertensive nephrosclerosis and fibrosis. It also demonstrated finerenone’s superior efficacy in reducing proteinuria and modulating these pathways (82). These findings, validated in human samples, support our study’s results, highlighting finerenone’s role in kidney protection and its therapeutic potential.

Diet induced obesity significantly contributes to mitochondrial dysfunction, characterized by impaired energy metabolism, altered mitochondrial morphology including changes in mitochondrial dynamics and reduced oxidative phosphorylation, all of which drive the progression of kidney disease (67–69). Studies in mice fed high-fat diet fed (HFD) and HFD with streptozotocin (HFD/STZ) induced diabetic mice (68, 83) have shown substantial production of reactive oxygen species (ROS) in the kidneys, accompanied by abnormal mitochondrial morphology, increased mitochondrial fission and mitophagy, particularly in proximal tubular cells (68, 83). At the molecular level, the oxidative stress marker Gp91 was markedly upregulated in the kidneys of HFD-fed mice, further highlighting the oxidative stress associated with obesity-induced kidney injury (68). In our study, a notable increase in the free-to-bound NADH ratio was observed in kidney sections from the western diet (WD) fed mice, reflecting increased glycolysis and reduced OXPHOS. This imbalance underscores the metabolic disruptions in mitochondrial energy pathways induced by diet-induced obesity. Importantly, finerenone treatment restored the mitochondrial metabolic balance in the kidneys of WD-fed mice, normalizing the free-to-bound NADH ratio. In addition, we have shown that finerenone treatment increased complex I activity in western diet fed mice. These findings suggest that finerenone effectively counteracts the mitochondrial dysfunction caused by diet-induced obesity, potentially restoring renal function.

To uncover the molecular mechanisms underlying the protective effects of finerenone, we investigated nuclear receptors and signaling pathways crucial for mitochondrial regulation. Among these, the estrogen-related receptor γ (ERRγ) is a key regulator of oxidative phosphorylation (OXPHOS) and fatty acid oxidation (84). While ERRγ and its isoform ERRα have been extensively studied in muscle and hepatic cells, their roles in renal mitochondria under metabolic stress remain less understood. Previous research demonstrated that the PGC-1α/ERRα axis regulates pyruvate dehydrogenase kinase 4 (PDK4) expression, reducing glucose oxidation rates in C2C12 myotubes (85). Additionally, ERRα was found to bind PDK1 and PDK2 in mouse livers, highlighting its role in modulating the metabolic switch between glucose and lipid utilization by mitochondria. Studies have identified estrogen-related receptors (ERRα, ERRβ, and ERRγ) as critical regulators of mitochondrial dysfunction and inflammation in the aging kidney (74). Notably, treatment with a pan-ERR agonist significantly reversed age-related renal damage, including albuminuria, mitochondrial impairment, and inflammation. In alignment with prior research, our study is the first to reveal that a western diet (WD) significantly reduces ERRγ expression in the kidneys, without any significant alterations in ERRα levels (74). Importantly, finerenone treatment restored ERRγ expression in WD-fed animals and increased ERRα expression, further supporting its role in enhancing mitochondrial function. These findings emphasize the critical involvement of ERRs in regulating mitochondrial energy metabolism in renal tissue under conditions of diet-induced obesity.

Given the established role of AMP-activated protein kinase (AMPK) in maintaining mitochondrial homeostasis and its impairment in renal dysfunction associated with obesity (86), we also evaluated AMPK signaling. WD-fed mice exhibited reduced phosphorylated AMPKα (p-AMPKα) levels relative to total AMPKα, indicative of compromised AMPK activity. Notably, finerenone treatment significantly restored p-AMPKα levels, suggesting that mineralocorticoid receptor inhibition reactivates AMPK signaling, promoting mitochondrial function and renal health. Reports have indicated that AMPK activation promotes ERRα expression and facilitates its binding to angiogenic gene promoters in muscle cells (87). Additionally, suppression of ERRα prevents the AMPK-induced upregulation of angiogenic factors, underscoring the pivotal role of ERRα in this regulatory pathway.

Importantly, obesity significantly contributes to chronic kidney disease (CKD) by promoting comorbid conditions like diabetes and hypertension and causing direct renal damage. This study highlights finerenone’s potential as a therapeutic agent, demonstrating its ability to mitigate obesity-induced renal injury through improved mitochondrial function, reduced inflammation, and fibrosis. By restoring key metabolic regulators like ERRγ and AMPK, finerenone preserves renal structure and function. Its efficacy in reducing markers of kidney injury and protecting podocytes underscores its promise in managing obesity-associated CKD. These findings provide a strong rationale for clinical translation to improve outcomes in CKD patients with metabolic dysfunction. In summary, our findings demonstrate that finerenone mitigates obesity-related renal damage by counteracting mitochondrial dysfunction. It enhances OXPHOS, preserves mitochondrial integrity, and re-establishes key regulatory pathways, including ERRγ and AMPK signaling. These results provide novel mechanistic insights into the renoprotective effects of finerenone and support its therapeutic potential in managing kidney disease associated with diet-induced obesity.

## SUPPLEMENTAL MATERIAL

Supplemental figures 1–2 and table 1: https://doi.org/10.6084/m9.figshare.29649716

