## supplemental file for "The non-steroidal MR antagonist Finerenone reverses Western diet-induced kidney disease by regulating mitochondrial and lipid metabolism and inflammation"

### Supplementary Figure 1: Measurement of body weight

**A**

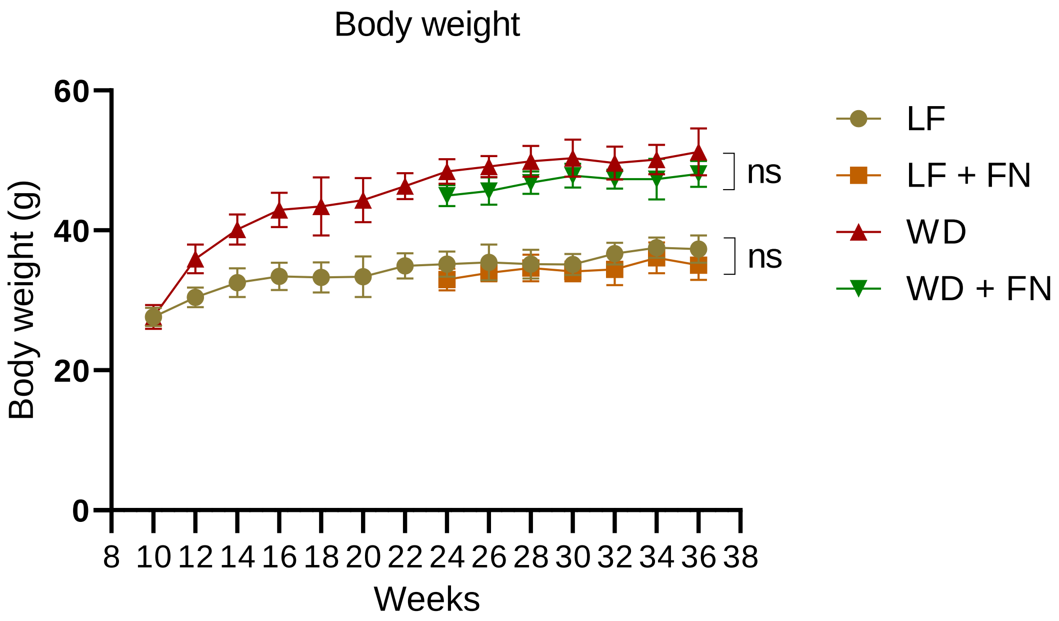

Supplementary Figure 1: Measurement of food intake

**B**

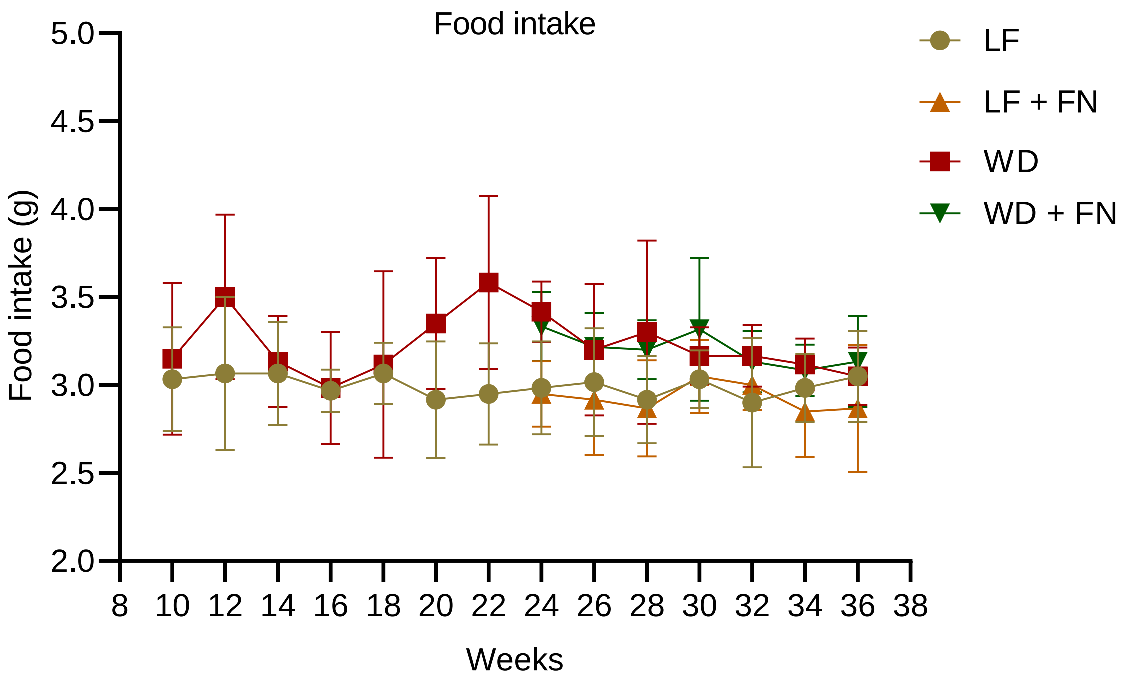

### Supplementary Table: List of primers

| Gene name | Forward | Reverse |
|---|---|---|
| **Fn1** | ATGTGGACCCCTCCTGATAGT | GCCCAGTGATTTCAGCAAAGG |
| **Col4a2** | GACCGAGTGCGGTTCAAAG | CGCAGGGCACATCCAACTT |
| **Pai1** | GACTCTGGATGAGTGGAAAGC | GGCGGCCTAAGTCTCCAAAAT |
| **TGFb** | CTCCCGTGGCTTCTAGTGC | GCCTTAGTTTGGACAGGATCTG |
| **CTGF** | GGGCCTCTTCTGCGATTTC | ATCCAGGCAAGTGCATTGGTA |
| **Timp1** | GCAACTCGGACCTGGTCATAA | CGGCCCGTGATGAGAAACT |
| **MCP1** | TTAAAAACCTGGATCGGAACCAA | GCATTAGCTTCAGATTTACGGGT |
| **NLRP3** | GCCTTGAAGAAGAGTGGATGG | GCGTGTAGCGACTGTTGAG |
| **cGAS** | GAGGCGCGGAAAGTCGTAA | TTGTCCGGTTCCTTCCTGGA |
| **TMEM173** | GGTCACCGCTCCAAATATGTAG | CAGTAGTCCAAGTTCGTGCGA |
| **TLR2** | GCAAACGCTGTTCTGCTCAG | AGGCGTCTCCCTCTATTGTATT |
